# Early-peaking caspase-7 activity at the plasma membrane drives apoptotic phosphatidylserine exposure

**DOI:** 10.1101/2025.03.17.643603

**Authors:** Yusuke Taira, Zhuohao Yang, Yugo Miyata, Yoshitaka Shirasaki, Katsumori Segawa, Masayuki Miura, Natsuki Shinoda

## Abstract

Apoptosis is an immunologically silent form of regulated cell death executed by caspase^1^. Caspase cleaves hundreds of substrates throughout the cell to regulate apoptotic processes^2^, including phosphatidylserine (PS) externalization^3–5^. However, spatio-temporal regulation of caspase activity in dying cells remains unclear. Here, we show that caspase activity peaks earlier at the plasma membrane (PM) during apoptosis by establishing a Dual Förster resonance energy transfer (Dual FRET) imaging system that combines subcellularly targeted FRET-based caspase biosensors with a cytosolic reference counterpart^6,7^. Genetic analysis identified caspase-7, an executioner caspase considered an inefficient backup for caspase-3, the major executioner caspase^8,9^, as a caspase responsible for the <u>E</u>arly-<u>P</u>eaking <u>C</u>aspase <u>A</u>ctivity at the <u>P</u>M (EP-CAP). Mechanistically, EP-CAP is mediated via electrostatic interactions between PS in the inner leaflet of the PM and polybasic residues in the N-terminal intrinsically disordered region (IDR) of caspase-7, which are liberated by the caspase-mediated removal of polyacidic residues. Physiologically, EP-CAP facilitates the efficient cleavage of phospholipid scramblases for the rapid externalization of PS and subsequent efferocytosis. Accordingly, we propose that caspase-7, but not caspase-3, is a bona fide immunologically silent death caspase reinforcing the non-inflammatory nature of apoptosis via EP-CAP.

## Main

Apoptosis is an immunologically silent form of regulated cell death that facilitates tissue homeostasis by eliminating unwanted cells from the body^1^. Apoptotic cells are rapidly eliminated by phagocytosis, termed efferocytosis^10,11^, the failure of which results in secondary necrosis that provokes inflammation, leading to several diseases, including atherosclerosis and systemic lupus erythematosus^12,13^. Caspase, a cysteine-aspartic acid protease, is known to be an executioner of apoptosis^1^. Upon apoptotic stimuli, initiator caspases, caspase-8 and caspase-9, are first activated and subsequently cleave downstream executioner caspases, caspase-3 and caspase-7^14,15^, where caspase-3 plays a major role, while caspase-7 is historically considered an inefficient backup^8,9^. Executioner caspases cleave hundreds of substrates throughout the cell to regulate corresponding apoptotic processes^2^. For example, executioner caspases cleave the inhibitor of caspase-activated DNase (ICAD) in the nucleus to activate CAD for DNA fragmentation^16–18^. For apoptotic cell clearance, executioner caspases cleave phospholipid translocases at the PM, such as Xkr8^3^, a phospholipid scramblase, and ATP11A/C^4,5^, phospholipid flippases, to facilitate PS exposure as an “eat-me” signal for efferocytosis^10,11^. Given the versatile roles of executioner caspases during apoptosis, their activities must be tightly regulated in a spatio-temporal order within the cell. This is further supported by the fact that caspase activity is spatio-temporally regulated in non-lethal scenarios^19–26^. However, owing to technical shortages, little is known regarding when, where, or even whether executioner caspase activity is subcellularly regulated within the cell during apoptosis.

### Executioner caspase activity peaks earlier at the PM during apoptosis

To visualize the spatio-temporal executioner caspase activity during apoptosis with subcellular resolution, we developed a Dual FRET imaging system (Fig.1 a-j) co-expressing subcellularly targeted O-DEVD-FR^7^, which is composed of mKOκ and mKate2 (Extended Data Fig. 1a, b), and cytosolic mSCAT3^6,25^, comprising mECFP and mVenus (Extended Data Fig. 1c, d), both of which can detect executioner caspase, namely caspase-3 and caspase-7, activity with a high temporal resolution during apoptosis by reducing the FRET ratio in HeLa cells. Accordingly, we selected O-DEVD-FR as a subcellularly targeted probe because O-DEVD-FR is more precisely localized to the destination of interest than mSCAT3 (Extended Data Fig. 1e–l), is cleaved upon apoptosis induction (Extended Data Fig. 1m–q), and that the localization of O-DEVD-FR is more stable than mSCAT3, as confirmed by fluorescence recovery after photobleaching (FRAP) analysis (Extended Data Fig. 2a–e), which is well-suited for analyzing subcellular caspase activity. We fused subcellularly targeted O-DEVD-FR and mSCAT3 via the P2A sequence to express equimolar amounts of both probes (Extended Data Fig. 3a–f) with independent subcellular localization (Extended Data Fig. 3g).

**Figure 1.**
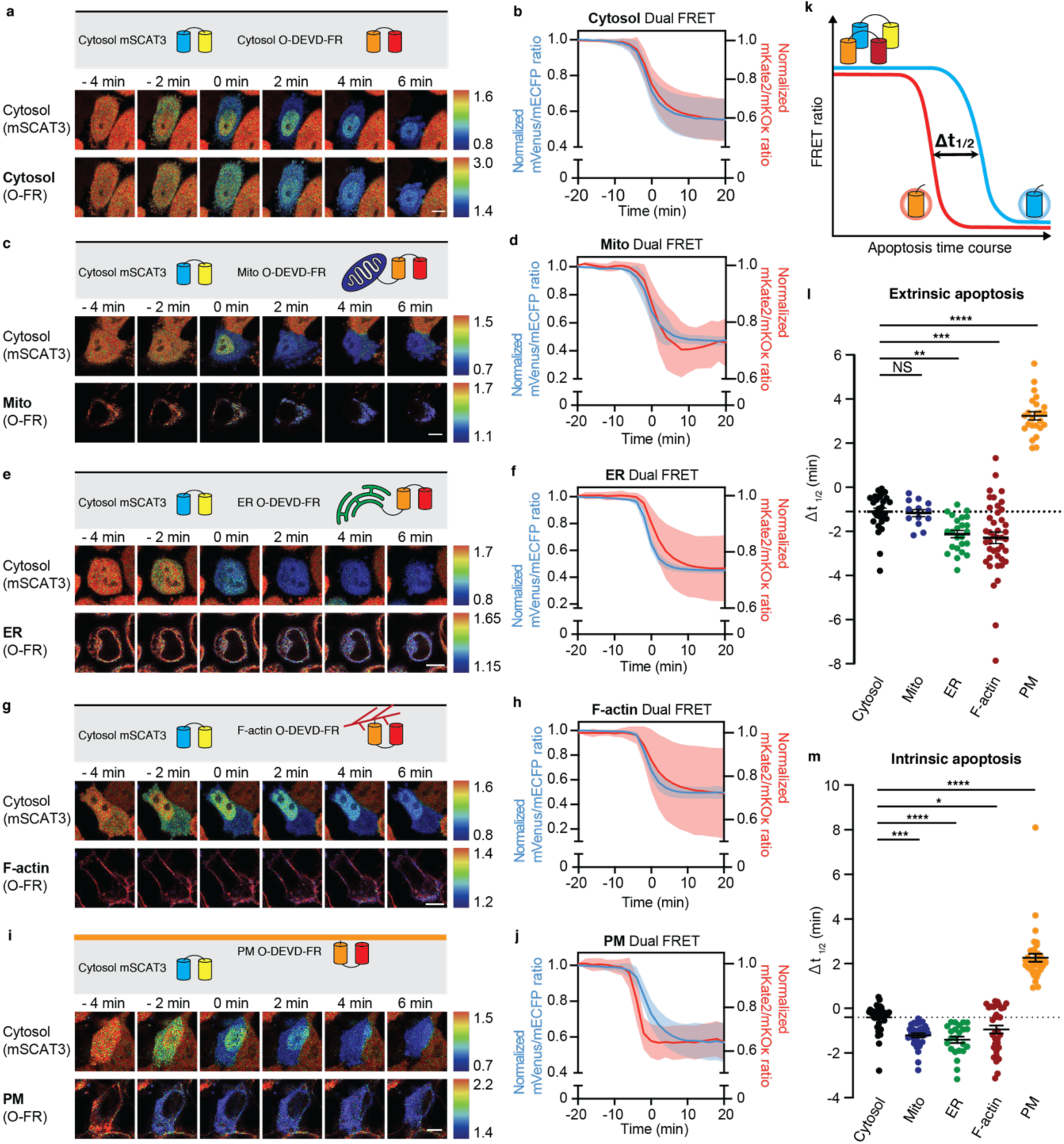
Dual FRET system identifies early-peaking caspase activity at the plasma membrane during apoptosis. (a-j) Live imaging of Dual FRET stably expressed in WT HeLa cells during extrinsic apoptosis at the indicated time. Zero min indicates the time when mSCAT3 FRET ratio reached half the maximum and minimum values. The FRET ratio is normalized by the ratio at t = -20 min. (a, c, e, g, i) Representative pseudo-colored FRET ratio images of cytosolic mSCAT3 (top: mVenus/mECFP) and subcellularly localized O-DEVD-FR (bottom: mKate2/mKOκ) in HeLa cells during extrinsic apoptosis. ((a) Cytosol, (c) mitochondria (Mito), (e) ER, (g) F-actin, (i) plasma membrane (PM)). Scale bars, 10 μm. (b, d, f, h, j) Average response of Dual FRET stably expressed in WT HeLa cells during extrinsic apoptosis (blue, cytosolic mSCAT3 vs red, subcellularly localized O-DEVD-FR). ((b) Cytosol, n = 30 cells; (d) Mito, n = 13 cells; (f) ER, n = 24 cells; (h) F-actin, n = 42 cells; (j) PM, n = 24 cells). (k) A schematic diagram illustrating the calculation of Δt_1/2_ in the Dual FRET system. Δt_1/2_ is measured to quantify the relative time of subcellularly localized caspase activation. (l) Δt_1/2_ of Cytosol Dual FRET (black, n = 30 cells (b)), Mito Dual FRET (blue, n = 13 cells (d)), ER Dual FRET (green, n = 24 cells (f)), F-actin Dual FRET (red, n = 42 cells (h)), and PM Dual FRET (orange, n = 24 cells (j)) during extrinsic apoptosis in WT HeLa. (m) Δt_1/2_ of Cytosol Dual FRET (black, n = 45 cells (Extended Data Fig. 4d)), Mito Dual FRET (blue, n = 31 cells (Extended Data Fig. 4f)), ER Dual FRET (green, n = 23 cells (Extended Data Fig. 4h)), F-actin Dual FRET (red, n = 33 cells (Extended Data Fig. 4j)), and PM Dual FRET (orange, n = 40 cells (Extended Data Fig. 4l)) during intrinsic apoptosis in WT HeLa. For all figures, time courses show the mean ± SD, and dot plots show the mean ± SEM. Statistical analysis was performed using one-way ANOVA with Dunnett’s multiple comparison test. NS: *P* > 0.05; *: *P* < 0.05; **: *P* < 0.01; ***: *P* < 0.001; ****: *P* < 0.0001.

We analyzed the caspase activity kinetics of the designated region by comparing the dynamics in FRET ratios of both probes, especially the time at which the ratio of each FRET reached half the maximum and minimum values (t_1/2 (mSCAT3)_- t_1/2 (O-DEVD-FR)_ (Δt_1/2_); Fig. 1k). During TNF-α and cycloheximide (CHX)-induced extrinsic apoptosis, the FRET ratio of cytosolic mSCAT3 and cytosolic O-DEVD-FR decreased at approximately similar rates (Δt_1/2_ = -1.10 ± 0.83 min; Fig. 1a, b, l), suggesting that O-DEVD-FR and mSCAT3 detect caspase activity with similar sensitivity. Caspase activity kinetics around mitochondria was similar to that in the cytosol (Δt_1/2_ = -1.17 ± 0.58 min; Fig. 1c, d, l). In contrast, caspase activity kinetics were delayed at both the endoplasmic reticulum (ER) and F-actin compared with those in the cytosol (ER: Δt_1/2_ = -2.12 ± 0.78 min; Fig. 1e, f, l), (F-actin: Δt_1/2_ = -2.29 ± 1.68 min; Fig. 1g, h, l). Conversely, caspase activity peaked significantly earlier at the PM than in the cytosol (Δt_1/2_ = 3.23 ± 0.90 min; Fig. 1i, j, l). Taken together, these results suggest that caspase activity is differentially regulated within cells during apoptosis, and caspase activity peaks earlier at the PM.

In mammals, apoptosis is triggered by either the intrinsic or extrinsic pathways, which are classified with the involvement of death receptors^14^. Next, we investigated the spatio-temporal profile of executioner caspase activity in the intrinsic pathway. There was no difference in the cytosolic mSCAT3 ratio kinetics and cleavage efficiency between extrinsic apoptosis and staurosporine (STS)-induced intrinsic apoptosis (Extended Data Fig. 4a, b). Additionally, there was almost no difference in the kinetics of FRET ratio reduction in cytosolic mSCAT3 and cytosolic O-DEVD-FR (Δt_1/2_ = -0.40 ± 0.56 min; Extended Data Fig. 4c, d, Fig. 1m). Similar to the extrinsic apoptosis pathway, we found that caspase activity was delayed around the mitochondria, ER, and F-actin (mitochondria: Δt_1/2_ = -1.21 ± 0.52 min, Extended Data Fig. 4e, f, Fig. 1m; ER: Δt_1/2_ = - 1.41 ± 0.71 min, Extended Data Fig. 4g, h, Fig. 1m; F-actin: Δt_1/2_ = -0.94 ± 1.02 min, Extended Data Fig. 4i, j, Fig. 1m). Furthermore, the executioner caspase activity peaked significantly earlier at the PM, even during intrinsic apoptosis (PM: Δt_1/2_ = 2.26 ± 1.13 min, Extended Data Fig. 4k, l, Fig. 1m). These results suggest that differentially regulated spatio-temporal caspase activity within the cell, including <u>E</u>arly <u>P</u>eaking executioner <u>C</u>aspase <u>A</u>ctivity at the <u>P</u>M (EP-CAP), is a universal feature of apoptosis, regardless of the apoptotic pathway.

### EP-CAP is mediated by caspase-7 with its IDR

Next, we examined the underlying molecular mechanisms of subcellularly regulated caspase activity, focusing on the PM. Given that EP-CAP is independent of apoptotic stimuli, we examined whether it is an emerging feature of the executioner caspases required for both pathways. Caspase-3 and caspase-7 both preferentially cleave DEVD sequences and exhibit similar substrate specificities^27^. However, recent studies revealed that their functions are not entirely identical^28–30^. To examine the contribution of each caspase to EP-CAP, we generated *caspase-3* knockout (KO) and *caspase-7* KO HeLa cells using CRISPR-Cas9 system (Extended Data Fig. 5a). We first investigated whether cytosolic Dual FRET kinetics differed among *caspase-3* KO, *caspase-7* KO, and wild-type (WT) cells. We found that *caspase-3* KO HeLa cells exhibited a slightly delayed decrease in the FRET ratio, possibly due to a delay in the overall apoptotic process, whereas *caspase-7* KO HeLa cells showed no significant difference compared with WT cells (Extended Data Fig. 5b–e). At the PM (WT PM Δt_1/2_ = 3.29 ± 1.45 min; Fig. 2a, d), activation kinetics differed in the established KO cells. Notably, EP-CAP was observed in *caspase-3* KO HeLa cells (*Caspase-3* KO PM Δt_1/2_ = 10.35 ± 3.54 min; Fig. 2b, d) but was depleted in *caspase-7* KO HeLa cells (*caspase-7* KO PM Δt_1/2_ = -2.00 ± 0.80 min; Fig. 2c, d), suggesting that EP-CAP is specifically mediated by caspase-7.

**Figure 2.**
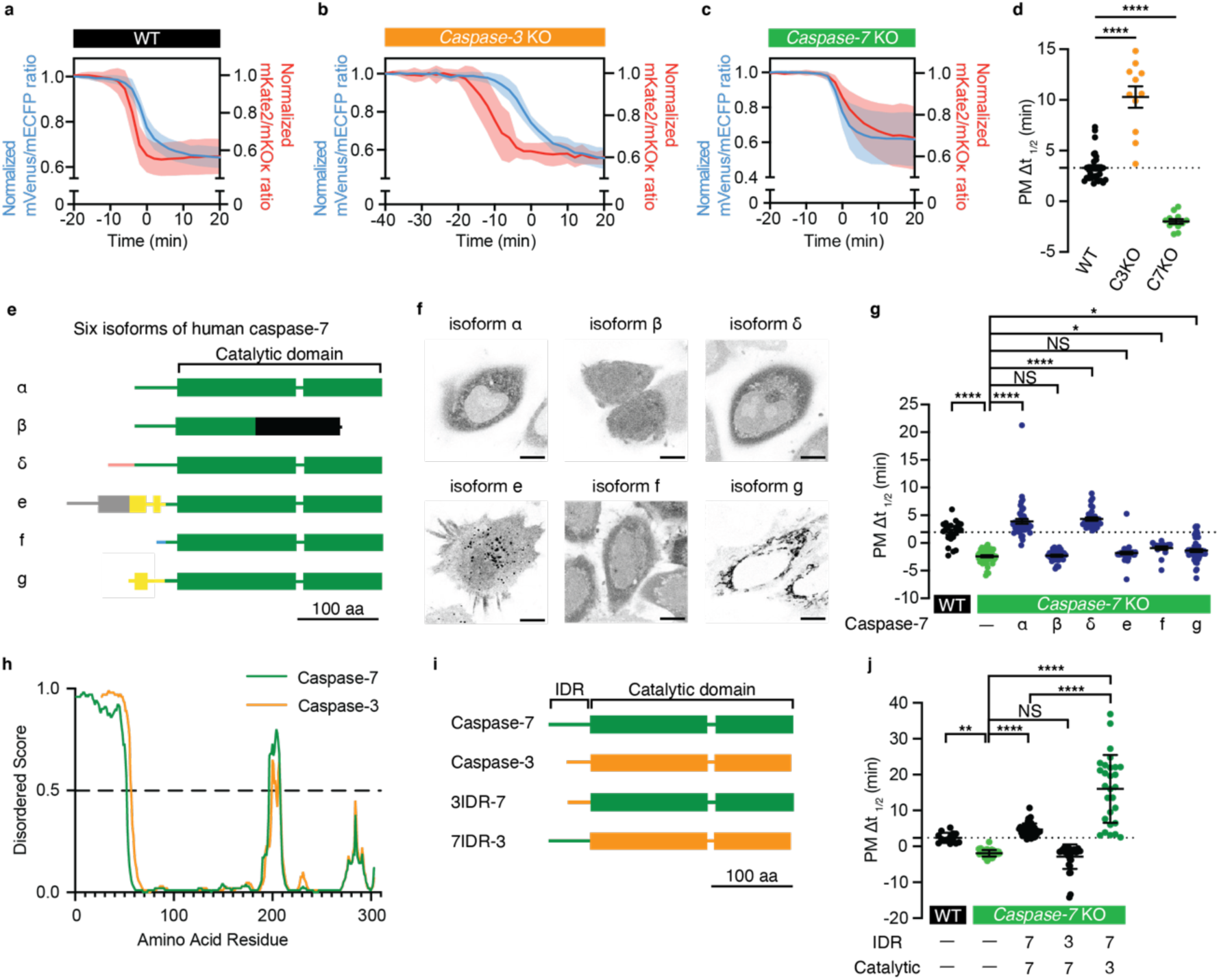
IDR of caspase-7 facilitates early-peaking caspase activity at the plasma membrane. (a-c) Average response of PM Dual FRET stably expressed in HeLa cells during extrinsic apoptosis (blue, cytosolic mSCAT3 vs red, PM O-DEVD-FR) (a, WT, n = 31 cells; b, *caspase-3* KO, n = 11 cells; c, *caspase-7* KO, n = 12 cells). The FRET ratio is normalized by dividing the ratio at t = -20 min (a and c) or t = -40 min (b). (d) Δt_1/2_ of PM Dual FRET in WT (black, n = 31 cells), *caspase-3* KO (orange, n = 11 cells), and *caspase-7* KO HeLa (green, n = 12 cells) during extrinsic apoptosis. (e) Schematic protein structures illustrating human caspase-7s. Isoform β and δ have the prodomain of isoform α (major isoform), but isoform β loses the catalytic domain (green boxes). Predicted intrinsically disordered regions (IDRs) are shown in lines. The same color lines indicate identical amino acid sequences, respectively. (f) Representative images of each caspase-7 isoform C-terminally fused with mNeonGreen expressed in HeLa cells. Scale bars, 10 μm. (g) Δt_1/2_ of PM Dual FRET transiently expressed in WT (n = 22 cells), *caspase-7* KO (n = 47 cells), *caspase-7* KO expressing caspase-7 isoform α-FLAG (n = 49 cells), caspase-7 isoform β-FLAG (n = 30 cells), caspase-7 isoform δ-FLAG (n = 31 cells), caspase-7 isoform e-FLAG (n = 38 cells), caspase-7 isoform f-FLAG (n = 18 cells), and caspase-7 isoform g-FLAG (n = 58 cells). Unless otherwise stated, caspase-7 refers to caspase-7 isoform α. (h) IDR predictions of caspase-3 (orange) and caspase-7 (green). Regions wherein the score is more than 0.5 are predicted to be disordered. Both caspases were described as C-terminus-aligned. (i) Schematic structures illustrating chimera caspase, in which IDR and catalytic domain are swapped. (j) Δt_1/2_ of PM Dual FRET transiently expressed in WT (n = 14 cells), *caspase-7* KO (n = 44 cells), *caspase-7* KO expressing caspase-7-FLAG (n = 40 cells), 3IDR-7-FLAG (n = 29 cells), 7IDR-3-FLAG (n = 24 cells). For all figures, time courses show the mean ± SD, and dot plots show the mean ± SEM. Statistical analysis was performed using one-way ANOVA with Dunnett’s multiple comparison test. NS: *P* > 0.05; *: *P* < 0.05; **: *P* < 0.01; ****: *P* < 0.0001.

Human caspase-7 encodes six protein isoforms, five of which possess identical catalytic domains, except for isoform β, whereas the N-terminal prodomain is distinct (Fig. 2e). Upon expressing mNeonGreen-tagged caspase-7 isoforms in HeLa cells, we found that caspase-7 isoform α, β, δ, and f diffuse throughout the cytosol, while caspase-7 isoform e and g exhibited puncta- and mitochondria-like structures, respectively (Fig. 2f), suggesting that the N-terminal prodomain regulates the localization of caspase-7. To identify isoforms required for EP-CAP, we conducted rescue experiments and found that only caspase-7 isoform α and δ recover EP-CAP during apoptosis; caspase-7 f did not demonstrate this activity, although it retains its catalytic domain and displays a localization pattern similar to α and δ (Fig. 2g). These results suggest that caspase-7, especially its N-terminal IDR composed of 1–52 aa in isoform α that covers isoform α and δ but not f, is responsible for EP-CAP during apoptosis (Fig. 2h).

To further validate whether the N-terminal IDR of caspase-7 contributes to EP-CAP, we generated chimeric caspases by swapping the IDR and catalytic domain in caspase-3 and caspase-7 (Fig. 2i). In *caspase-7* KO cells, caspase-3 IDR fused to the caspase-7 catalytic domain (3IDR-7) failed to rescue EP-CAP. In contrast, caspase-7 IDR fused with caspase-3 catalytic domain (7IDR-3) dramatically enhanced EP-CAP (3IDR-7 PM Δt_1/2_ = -1.83 ± 2.00 min, 7IDR-3 PM Δt_1/2_ = 22.07 ± 3.25 min; Fig. 2j). These results indicate that for caspase-7, rather than the catalytic domain, IDR in the prodomain is required and sufficient for EP-CAP.

### Cleavage of caspase-7 IDR gates EP-CAP

The caspase catalytic domain is necessary for cleaving substrates; however, there are few reports on the function of the caspase-7 IDR. IDR mediates weak multivalent molecular interactions and facilitates liquid-liquid phase separation^31^. However, no lipid droplet-like structures were observed in cells transiently overexpressing mNeonGreen-tagged caspase-7 (Fig. 2f). Interestingly, we noted that caspase-7 IDR possesses three unique features: a polyacidic motif, a polybasic motif, which reportedly interacts with RNA^32,33^, and a caspase cleavage site in between (Fig. 3a). These three characteristic patterns are well conserved in vertebrates, but not in invertebrates (Extended Data Fig. 6). Additionally, the IDR of caspase-7 was predicted to bind to the membrane using the basic and hydrophobic (BH) search tool^34^ (Fig. 3b, c). This region is present only in caspase-7 isoform α and δ but not in other isoforms of caspase-7 or in caspase-3. To determine the importance of charged regions in EP-CAP, we mutated nine basic amino acids [lysine (K) and arginine (R)] in the polybasic motif (KR/A), and seven acidic amino acids [aspartic acid (D) and glutamic acid (E)] in the polyacidic motif (ED/A) (Fig. 3a). To specifically evaluate the function of the IDR in caspase-7, we performed PM Dual FRET analysis using 7IDR-3, in which the catalytic domain was derived from caspase-3. Using charged amino acid-mutated chimeric caspases, we demonstrated that in *caspase-7* KO cells, 7IDR^KR/A^-3 did not recover the delayed caspase activity at the PM, whereas 7IDR^ED/A^-3 significantly promoted EP-CAP (Fig. 3d), suggesting that basic amino acids are required for EP-CAP, while acidic amino acids suppress it. Next, we evaluated the effect of proteolytic cleavage of this sequence by mutating the cleavage site in caspase-7 IDR (D23A) (Fig. 3a). Interestingly, 7IDR^D23A^-3 did not rescue the delayed caspase activity at the PM (Fig. 3e). Taken together, our results suggest that the clustered polybasic region, liberated by the proteolytic removal of clustered polyacidic region, are crucial for EP-CAP.

**Figure 3.**
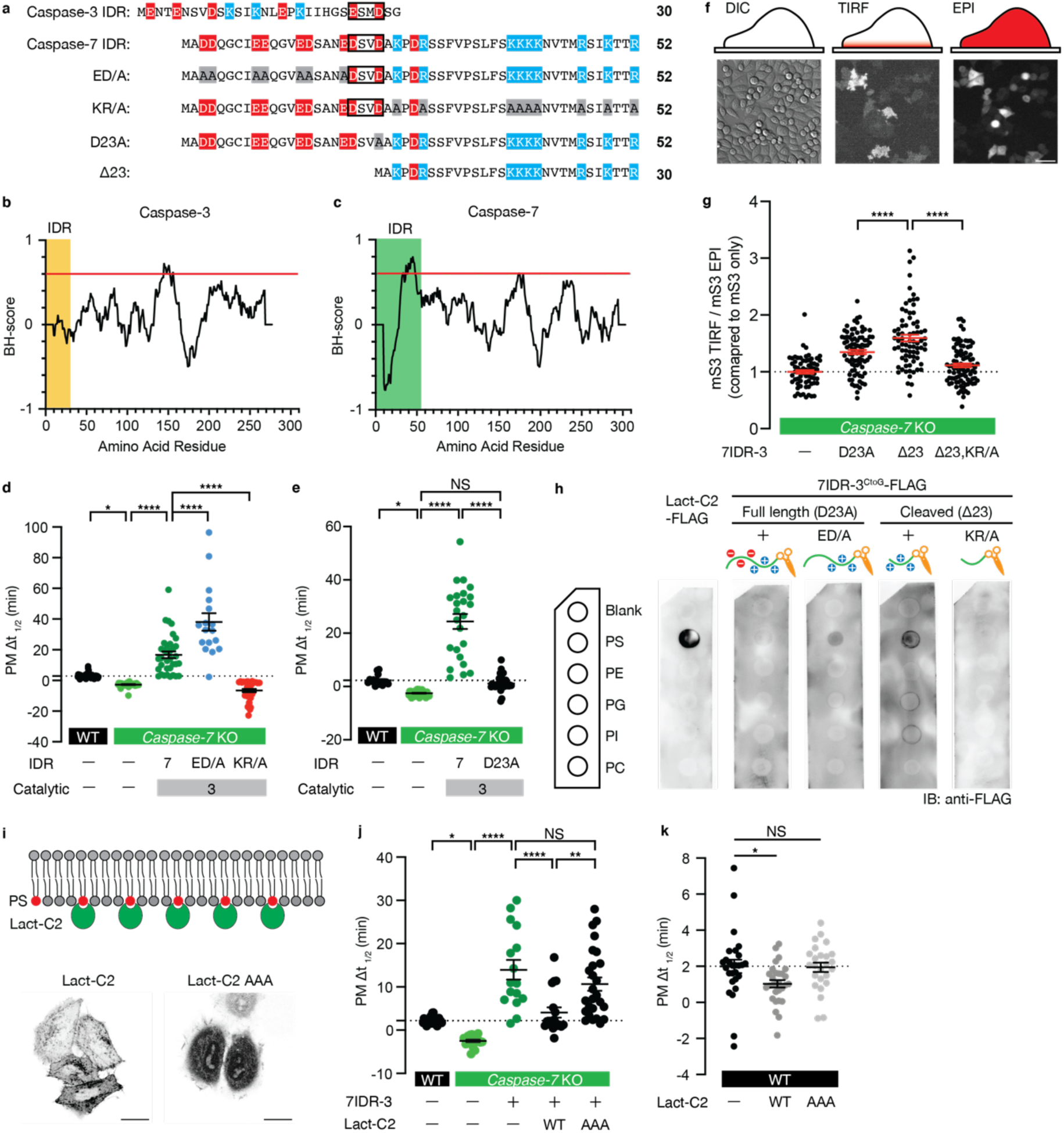
Caspase-7-PS electrostatic interaction gated by proteolytic cleavage is necessary for early-peaking caspase activity at the plasma membrane. (a) Alignment of the amino acid sequence of caspase-3 IDR, caspase-7 IDR, IDR mutants. Acidic amino acids are shown in red. Basic amino acids are shown in blue. Caspase recognition sites are shown in boxes. Mutated amino acids are highlighted in gray. (b, c) Basic and hydrophobic (BH)-scores of caspase-3 and caspase-7. The red line indicates a threshold of 0.6. Caspase-3 IDR and caspase-7 IDR are masked with orange and green blocks, respectively. (d) Δt_1/2_ of PM Dual FRET transiently expressed in WT (n = 45 cells), *caspase-7* KO (n = 37 cells), with 7IDR-3-FLAG (n = 35 cells), 7IDR^ED/A^-3-FLAG (n = 17 cells), 7IDR^KR/A^-3-FLAG (n = 34 cells). (e) Δt_1/2_ of PM Dual FRET transiently expressed in WT (n = 15 cells), *caspase-7* KO (n = 44 cells), with 7IDR-3-FLAG (n = 24 cells), 7IDR^D23A^-3-FLAG (n = 31 cells). (f) A schematic image illustrating differential interference contrast (DIC), TIRF, and epifluorescence observation (top). Representative images of mScarlet3 expressing HeLa cells (bottom). Scale bar: 50 μm. (g) The ratio of TIRF fluorescence intensity to epifluorescence intensity, normalized to mScarlet3 (mS3) alone. (h) Lipid overlay assays with recombinant Lact-C2-FLAG, 7IDR^D23A^-3^CtoG^-FLAG, 7IDR^D23A, ED/A^- 3^CtoG^-FLAG, 7IDR^Δ23^-3^CtoG^-FLAG, and 7IDR^Δ23, KR/A^-3^CtoG^-FLAG. (i) A schematic diagram depicting Lact-C2 interacting with phosphatidylserine (PS) in the inner leaflet of the PM (top). Representative images of Lact-C2 fused with HaloTag7 (bottom). Scale bars: 10 μm. (j) Δt_1/2_ of PM Dual FRET transiently expressed in WT (n = 27 cells), *caspase-7* KO (n = 19 cells), with 7IDR-3-FLAG (n = 16 cells), 7IDR-3-FLAG + HaloTag7-Lact-C2 (n = 18 cells), 7IDR-3-FLAG + HaloTag7-Lact-C2^AAA^ (n = 26 cells). (k) Δt_1/2_ of PM Dual FRET transiently expressed in WT (n = 27 cells), with HaloTag7-Lact-C2 (n = 29 cells), HaloTag7-Lact-C2^AAA^ (n = 25 cells). For all figures, dot plots show the mean ± SEM. Statistical analysis was performed using one-way ANOVA with Dunnett’s comparison test. NS: *P* > 0.05; *: *P* < 0.05; **: *P* < 0.01; ****: *P* < 0.0001.

We then examined the changes in caspase-7 localization during apoptosis. However, we did not observe clear PM localization of caspase-7 even during apoptosis (Extended Data Fig. 7a, b). We speculate that proteolytic cleavage of IDR weakly increases the proportion of the PM-associated pool of caspase-7 for EP-CAP, which is undetectable by conventional confocal microscopy. To evaluate the proportion of caspases in the vicinity of the PM, we used total internal reflection fluorescence (TIRF), which enables the precise quantification of the PM-associated caspase-7 pool in terms of area and epifluorescence (Epi) microscopy (Fig. 3f). Compared with full-length (7IDR^D23A^-3), polyacidic regions-removed form (7IDR^Δ23^-3) increased the proportion of caspase localized to the PM (Fig. 3g). Moreover, the increased proportion of PM-localized caspase was abrogated when basic amino acids were mutated (7IDR^Δ23, KR/A^-3) (Fig. 3g). These results suggest that the polybasic region could enhance the proportion of caspase-7 associated with PM.

### Caspase-7 IDR interacts with PS upon cleavage for EP-CAP

Cellular membranes have distinct lipid compositions and a wide range of charges. PS is an anionic phospholipid that is abundant in mammalian PM^35^ and contributes substantially to the negative charge of the membrane^36^. Polybasic motif has a high affinity for anionic membrane lipids^37^. Accordingly, we examined the interaction between the caspase-7 IDR and phospholipids, including PS. To specifically examine the modification of caspase-7 IDR, we prepared full-length (D23A) and cleaved from (Δ23) of FLAG-tagged recombinant 7IDR-3 with caspase catalytically inactive (CtoG) to suppress self-cleavage (Extended Data Fig. 8). We found that the cleaved form (7IDR^Δ23^- 3^CtoG^) interacted specifically with PS, with the interaction dependent on polybasic amino acids (Fig. 3h). In contrast, the full-length form (7IDR^D23A^-3^CtoG^) did not interact with any phospholipids, whereas 7IDR^D23A^-3^CtoG^ carrying mutations in polyacidic residues (7IDR^D23A, ED/A^-3^CtoG^) bound to PS, suggesting that the proteolytic removal of polyacidic regions is essential for the interaction between caspase-7 IDR and PS (Fig. 3h). These results suggest that caspase-7 directly interacts with PS in the inner leaflet of the PM, with the timing regulated by apoptosis stimuli-induced cleavage of its IDR.

To examine the contribution of PS to EP-CAP, we masked PS headgroups by overexpressing the lactadherin C2 domain (Lact-C2)^36^ (Fig. 3i). Lact-C2 inhibited the EP-CAP induced by 7IDR-3 expression, whereas Lact-C2^AAA^, a mutant unable to bind PS, did not induce this inhibition (Fig. 3j). These results suggest that the polybasic residues in caspase-7 IDR interact with PS in the PM inner leaflet to facilitate EP-CAP. Furthermore, Lact-C2 expression delayed EP-CAP in WT HeLa cells (Fig. 3k), suggesting that PS in the inner leaflet of the PM is required for endogenous EP-CAP. Collectively, our results indicate that the electrostatic interaction between caspase-7 and PS in the inner leaflet of the PM, which is gated by caspase-mediated cleavage of IDR, facilitates timely and spatially regulated subcellular caspase activity during apoptosis.

### EP-CAP promotes apoptotic PS exposure and efferocytosis

Caspases execute apoptosis by cleaving diverse intracellular substrates. To determine the physiological function of EP-CAP, we examined its contribution to apoptotic PS exposure mediated by phospholipid scramblase Xkr8, whose activation is dependent on caspase-mediated cleavage^3^. Xkr8 has a caspase recognition site at its cytosolic C-terminus. Thus, we performed live imaging of HeLa cells expressing Xkr8 with EGFP fused to the C-terminus (Fig. 4a). At the onset of caspase activation, the EGFP signal was observed along the edge of the PM blebs; however, after 60 min, the signal diffused into the cytosol (Fig. 4b). Little cleavage-resistant (2DA) Xkr8-EGFP was detached from the PM, suggesting that EGFP detachment reflects the cleavage of Xkr8 (Fig. 4c). *Caspase-7* KO cells exhibited a slower rate EGFP signal decline at the PM after caspase activation than WT cells, suggesting that caspase-7 efficiently cleaves PM substrates (Fig. 4c). By establishing caspase-7 and IDR-modified caspase-7s stably expressing cells in *caspase-7* KO cells, which reproduced EP-CAP (Extended Data Fig. 9 a, b), we demonstrated that the reduced cleavage efficiency of Xkr8 was rescued by expressing caspase-7 but not by 3IDR-7 (Fig. 4c’). Among the 7IDR-3 variants, 7IDR-3 and 7IDR^ED/A^-3, which exhibited EP-CAP, rescued the reduced cleavage efficiency of Xkr8, whereas 7IDR^D23A^-3 and 7IDR^KR/A^-3 did not demonstrate this activity (Fig. 4c”). These results suggest that EP-CAP facilitates endogenous substrate cleavage at the PM.

**Figure 4.**
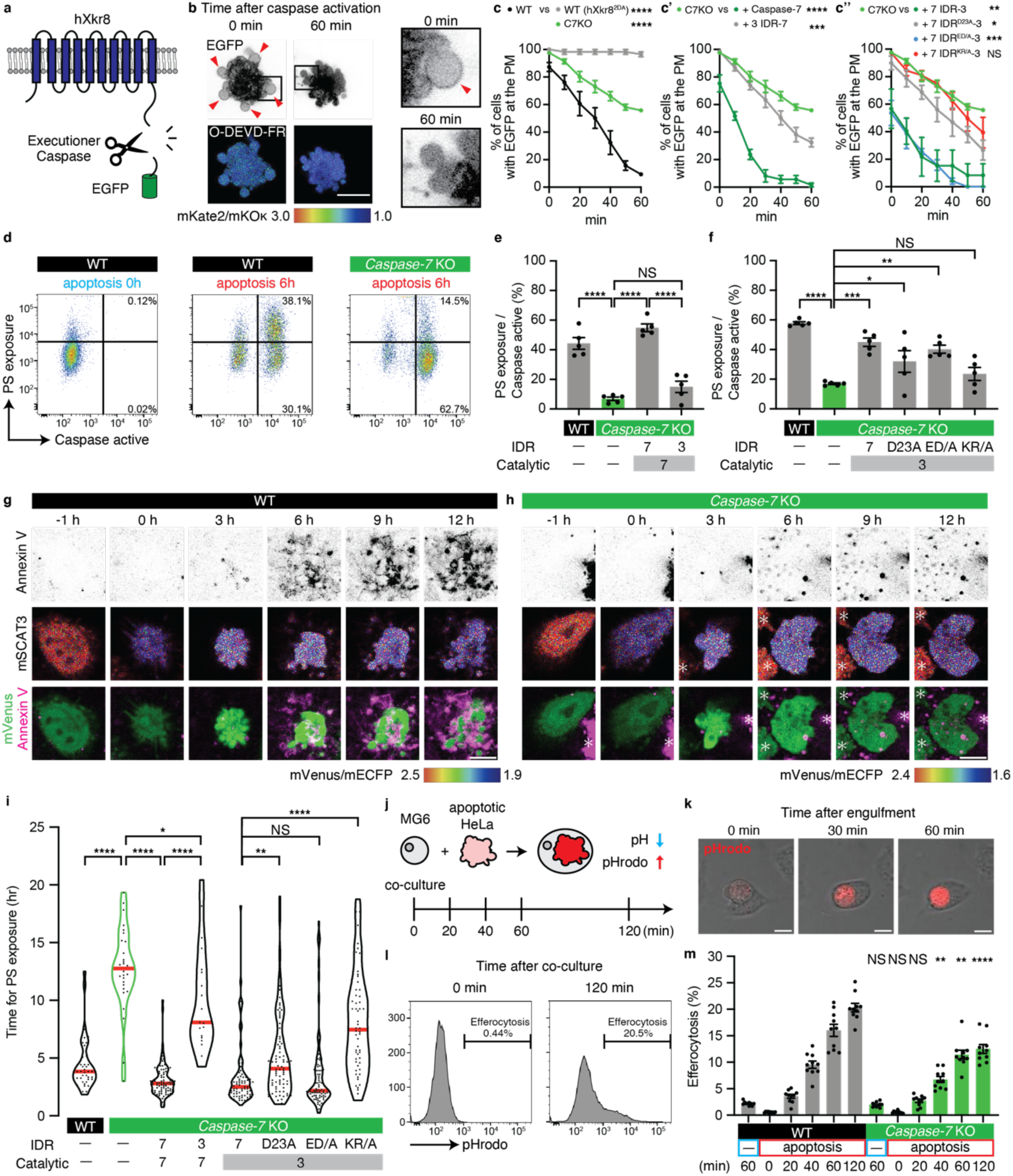
Early-peaking caspase activity at the plasma membrane facilitates rapid PS exposure and efferocytosis. (a) A schematic diagram illustrating hXkr8 with EGFP fused to the C-terminus. (b) Representative images of hXkr8-EGFP (top) and pseudo-colored FRET ratio images of cytosolic O-DEVD-FR (bottom: mKate2/mKOκ) in HeLa cells during extrinsic apoptosis at the indicated time. Red arrowheads indicate the EGFP signal remaining along the PM. Enlarged images of the boxed areas are shown on the right. (c, c’, c’’) Percentage of cells expressing hXkr8-EGFP, in which EGFP remains in the PM after caspase activation. Zero min indicates the time at which O-DEVD-FR ratio reaches its minimum. The experiments were independently repeated three times. The *caspase-7* KO curves shown in c’ and c” were reproduced from c. Statistical analysis was performed on data obtained 60 min after caspase activation. (d) Caspase activation (CellEvent) and PS exposure (APC-Annexin V) in HeLa cells with or without extrinsic apoptosis, as analyzed by flow cytometry. (e, f) The ratio of PS-exposing cells to caspase-active cells was analyzed by flow cytometry. The experiments were independently repeated five times. (g, h) Representative images of PS exposure in WT and *caspase-7* KO HeLa cells during extrinsic apoptosis. Zero hours indicates the time at which the mVenus/mECFP ratio reaches its minimum. Asterisks indicate neighboring cells. Scale bars, 10 μm. (i) Time from caspase activation to PS exposure in WT (n = 38 cells), *caspase-7* KO (n = 28 cells), with caspase-7-FLAG (n = 86 cells), 3IDR-7-FLAG (n = 19 cells), 7IDR-3-FLAG (n = 79 cells), 7IDR^D23A^-3-FLAG (n = 79 cells), 7IDR^ED/A^-3-FLAG (n = 79 cells), and 7IDR^KR/A^-3-FLAG (n = 56 cells) HeLa cells during extrinsic apoptosis. (j) A schematic diagram illustrating the efferocytosis assay using MG6 and pHrodo-labeled apoptotic HeLa cells. (k) Representative images showing MG6 engulfing apoptotic HeLa cells labeled with pHrodo. Scale bars, 10 μm. (l) Flow cytometry histogram showing the population of MG6 cells after co-culture with apoptotic HeLa cells labeled with pHrodo. (m) The percentage of MG6 engulfing apoptotic HeLa cells was analyzed by flow cytometry. WT and *caspase-7* KO under the same conditions were subjected to statistical analyses. For all figures, dot plots show the mean ± SEM. Statistical analysis was performed using one-way ANOVA with Dunnett’s comparison test (**c**, **e**, **f**, **i**) and unpaired two-tailed *t*-test (**m**). NS: *P* > 0.05; *: *P* < 0.05; **: *P* < 0.01; ***: *P* < 0.001; ****: *P* < 0.0001.

Next, we examined the physiological outcomes of the EP-CAP in apoptotic cells. Using flow cytometry analysis, we demonstrated that some caspase-activated cells exposed PS in the WT condition 6 h after apoptosis induction (Fig. 4d-f). In *caspase-7* KO cells, the ratio of PS-exposing cells to caspase-activated cells was reduced, suggesting that caspase-7 was required for rapid PS exposure during apoptosis (Fig. 4d-f). Caspase-7, but not 3IDR-7, expression rescued reduced PS exposure (Fig. 4e). Furthermore, reduced PS exposure was rescued by 7IDR-3 and 7IDR^ED/A^-3, only partially by 7IDR^D23A^-3 and not rescued by 7IDR^KR/A^-3 (Fig. 4f). These results support the hypothesis that polybasic residues in the caspase-7 IDR, which are absent in the caspase-3 IDR, are required for rapid PS exposure during apoptosis. We further investigated the kinetics of PS exposure at the single-cell level. We performed time-lapse imaging of PS exposure with caspase activity, followed by apoptosis induction using mSCAT3 and Annexin V. In this experimental system, WT cells required an average of 4.41 h to show PS exposure after caspase activation (Fig. 4g, i). *Caspase-7* KO cells exhibited delayed PS exposure (12.7 h) (Fig. 4h, i), suggesting that caspase-7 is required for rapid PS exposure during apoptosis. Caspase-7, but not by 3IDR-7, expression significantly rescued the delay in PS exposure in *caspase-7* KO cells (Fig. 4i). Among the mutant 7IDR-3s, rapid PS exposure was achieved by 7IDR-3 and 7IDR^ED/A^-3 but only weakly by 7IDR^D23A^-3 and not by 7IDR^KR/A^-3 (Fig. 4i), suggesting that EP-CAP facilitates rapid PS exposure. Finally, we evaluated efferocytotic efficiency by co-culturing apoptotic HeLa cells and the murine microglial cell line MG6 (Fig. 4j, k). We found that *caspase-7* KO cells were more resistant to efferocytosis mediated by MG6 than WT cells (Fig. 4l, m). Overall, these results suggest that EP-CAP facilitates rapid PS exposure and the subsequent efferocytosis of apoptotic cells (Extended Data Fig. 10).

## Discussion

Recent advances in subcellularly targeted biosensors have revealed spatial heterogeneity in the dynamics of AMPK^38^, ERK^39^, and PKC^40^. Similarly, caspases exhibit spatially restricted activity in non-lethal contexts to apoptosis^19–26^. However, given the limitations of existing probes, the spatio-temporal dynamics of subcellular caspase activity during apoptosis remain underexplored^6,7,24,41–43^. Here, using our Dual FRET imaging system, we revealed that caspases exhibit spatially heterogeneous activity patterns during apoptosis, with EP-CAP facilitating rapid “eat-me” signal exposure and efferocytosis. These findings suggest that caspase activity remains tightly regulated even under lethal conditions to control post-mortem cell fate.

Caspase-3 and caspase-7 share 54% amino acid sequence identity^44^ and preferentially cleave DEVD sequences^27^, although they exhibit substrate specificity *in vitro*. For example, Gasdermin E (GSDME), which is essential for pyroptosis and secondary necrosis via PM pore formation, is cleaved specifically by caspase-3^45–47^. In contrast, caspase-7 preferentially cleaves PARP-1 and other RNA-binding proteins^32,33^. Although both caspase-3 and caspase-7 cleave Xkr8 *in vitro*^3^, our *in-cellulo* results reveal that caspase-7 is preferentially required for Xkr8 cleavage during apoptosis. This substrate preference, driven by enzyme-lipid interactions, underscores the importance of analyzing cleavage kinetics directly in a spatio-temporal manner in complex cellular environments.

Charge blocks within proteins are determinants of subcellular localization, with IDRs driving liquid-liquid phase separation^48–50^ and polybasic motifs facilitating PM localization via electrostatic interactions^51,52^. Using the Dual FRET imaging system, we demonstrated that proteolytic cleavage of the IDR defines the spatial and temporal pattern of activity, acting as a molecular “ticket” that allows entry into the playground. This suggests that in addition to primary sequences, post-translational modifications that confer new properties should also be considered in regulatory analyses. Given that efferocytosis of infected apoptotic cells by dendritic cells facilitates antigen cross-presentation^53^, vertebrate-specific charge block patterns in caspase-7 IDR can contribute to adaptive immunity through EP-CAP. Furthermore, the predominant cytosolic localization of caspase-7 during apoptosis, with limited PM preference, suggests a regulatory mechanism that balances access to PM substrates with intracellular targets. This principle is particularly relevant for multitargeting enzymes such as caspases, highlighting the importance of *in-cellulo* analysis.

Given its EP-CAP, caspase-7 could also efficiently cleave other PM substrates, such as PS flippases, ATP11A/C^4,5^, which is inactivated by cleavages, and pannexin-1, which releases ATP as a “find-me” signal^54,55^ upon cleavage, further ensuring the non-inflammatory nature of apoptosis. Furthermore, GSDME, an executioner of secondary necrosis, is activated by caspase-3 but not by caspase-7^45–47^. In contrast, caspase-7 facilitates membrane repair by activating acid sphingomyelinase, thereby sealing GSDM-induced pores in mice^30^. Based on these findings, we propose that caspase-7 is an immunologically silent death caspase that reinforces the non-inflammatory nature of apoptosis.

## Methods

### Molecular cloning

The establishment of *pcDNA3.1-myc-mSCAT3*^25^ was previously reported. For *pcDNA3.1-V5-O-DEVD-FR*, the coding sequence of O-DEVD-FR was PCR-amplified using V5-tag harboring the primer from *pcDNA3.1-O-DEVD-FR* (a gift from H. Imamura)^7^. For *pcDNA3.1-V5-O-DEVG-FR*, the coding sequence of O-DEVG-FR was PCR-amplified using mutation-introducing (D238G, GAC to GGT) primer from *pcDNA3.1-V5-O-DEVD-FR*. For mitochondria-localized probes, the coding sequence of TOM20_1-36_ was PCR-amplified from the coding sequence corresponding to amino acids 1–36 of *Drosophila* TOM20 (NCBI Reference Sequence: NP_649169.3)^56^. For ER-localized probes, the coding sequence of Sec63_1-220_ was PCR-amplified from the coding sequence corresponding to amino acids 1–220 of *Drosophila* Sec63 (NCBI Reference Sequence: NP_648111.1)^56^. For F-actin-localized probes, the gene fragment of F-tractin^57,58^ was synthesized using Integrated DNA Technologies. For plasma membrane (PM)-localized probes, the coding sequences of myc-mSCAT3^DEVD^ and V5-O-DEVD-FR were PCR-amplified using a myristoylation sequence derived from avian Src (N-MGSSKSKPK-C)^59,60^ harboring primers for the coding sequence of myc-mSCAT3^DEVD^ and V5-O-DEVD-FR. The coding sequences of five types of subcellularly localized (cytosol, mitochondria, ER, F-actin, and PM) myc-mSCAT3 were ligated to the *Bam*HⅠ/*Hin*dⅢ-digested pcDNA3.1 vector using NEbuilder (#E2621L; New England Biolabs) DNA Assembly. The coding sequences of five variants of subcellularly localized V5-O-DEVD-FR were ligated to the *Bam*HⅠ/*Hin*dⅢ-digested pcDNA3.1 vector using In-Fusion (#639650; Takara).

For the five variants of V5-O-DEVD-FR-P2A-myc-mSCAT3^DEVD^, the coding sequences of V5-O-DEVD-FR and myc-mSCAT3^DEVD^ were PCR-amplified using the N-GGGSGGG-C linker and 2A peptide derived from porcine teschovirus-1^61^ (P2A, N-ATNFSLLKQAGDVEENPGP-C), harboring primers from pcDNA3.1-subcellularly localized V5-O-DEVD-FR and pcDNA3.1-myc-mSCAT3^DEVD^, respectively. The coding sequences were ligated to the *Bam*HⅠ/*Hin*dⅢ-digested pcDNA3.1 vector or the *Not*Ⅰ-digested *pPB-Gα15-FLAG-IRES-puroR* vector (a gift from A. Koto) ^62^ using NEbuilder HiFi DNA Assembly or In-Fusion, respectively.

For caspase-7 isoforms (α, β) of *pcDNA3.1-caspase-7s-FLAG*, the coding sequences of human caspase-7s were PCR-amplified using FLAG-tag harboring primers from human cDNA extracted from HeLa cells and were ligated to the *Bam*HⅠ/*Hin*dⅢ-digested pcDNA3.1 vector using In-Fusion. For caspase-7 isoforms (δ, e, f, g) of *pcDNA3.1-caspase-7s-FLAG*, the coding sequences of human caspase-7s were PCR-amplified using IDR harboring primers from *pcDNA3.1-caspase-7 α-FLAG* and were ligated to the *Bam*HⅠ/*Hin*dⅢ-digested pcDNA3.1 vector using In-Fusion. Unless otherwise stated, caspase-7 refers to caspase-7 isoform α.

For *pcDNA3.1-caspase-7s-myc-mNeonGreen*, the coding sequence of myc-mNeonGreen was PCR-amplified from *pUASz-Drice-myc-mNeonGreen*^24^ and ligated the coding sequence of human caspase-7s to the *Bam*HⅠ/*Hin*dⅢ-digested pcDNA3.1 vector using In-Fusion.

For *pcDNA3.1-3IDR-7-FLAG* and *pcDNA3.1-7IDR-3-FLAG*, the coding sequence of caspase-3 IDR (caspase-3 amino acids 1–30), caspase-3 catalytic domain (caspase-3 amino acids 31–277), caspase-7 IDR (caspase-7 amino acids 1–52), and caspase-7 catalytic domain (caspase-7 amino acids 53–303) were PCR-amplified using FLAG-tag harboring primers from *pcDNA3.1-caspase-7-FLAG* or *pcDNA3.1-caspase-3-FLAG*, and the two fragments of IDR/catalytic domain pairs derived from each caspase were ligated to the *Bam*HⅠ/*Hin*dⅢ-digested pcDNA3.1 vector using In-Fusion.

For mutant 7IDR-3-FLAG (ED/A, KR/A, D23A (GAT to GCC)), the coding sequence of caspase-7 IDRs was PCR-amplified using mutation-introducing primers from *pcDNA3.1-7IDR-3-FLAG* and ligated to the coding sequence of the caspase-3 catalytic domain-FLAG and the *Bam*HⅠ/*Hin*dⅢ-digested pcDNA3.1 vector using In-Fusion.

For the six variants of *pMXs-puro-caspase-FLAG* (caspase-7, 3IDR-7, 7IDR-3, 7IDR^ED/A^-3, 7IDR^KR/A^-3, and 7IDR^D23A^-3), the caspase coding sequences were PCR-amplified from *pcDNA3.1-caspase-FLAGs* and ligated to the *Bam*HⅠ/*Eco*RⅠ-digested pMXs-puro^63^ (a gift from T. Kitamura) vector using In-Fusion.

For *pcDNA3.1-HaloTag7-Lact-C2* and *pcDNA3.1-HaloTag7-Lact-C2^AAA^*, the coding sequence of Lact-C2 was PCR-amplified using primers for Lact-C2 or mutation-introducing primers (W26A; TGG to GCC, W33A; TGG to GCC, F34A; TTT to GCC) from *Lact-C2-GFP* (#22852; Addgene). The coding sequence of HaloTag7 was PCR-amplified using a HaloTag-PSMD3 fusion vector (#N2701; Promega). The two fragments were ligated to the *Bam*HⅠ/*Hin*dⅢ-digested pcDNA3.1 using In-Fusion.

For *pEF-EX-BOS-Lact-C2-FLAG*, the coding sequence of Lact-C2 was PCR-amplified using FLAG-tag harboring primers from *pcDNA3.1-HaloTag7-Lact-C2* and ligated to the *Pac*Ⅰ/*Bam*HⅠ-digested *pEF-EX-BOS vector*^64^ using In-Fusion. For *pEF-EX-BOS-Caspase-3^CtoG^-FLAG*, the coding sequence of caspase-3^CtoG^ was PCR-amplified using mutation-introducing (C163G)^65^ and FLAG-tag harboring primers from the coding sequence of caspase-3 and ligated to the *Pac*Ⅰ/*Not*Ⅰ-digested *pEF-EX-BOS* vector using In-Fusion.

For the four variants of *pEF-EX-BOS-7IDR-3^CtoG^-FLAG*, the catalytic domain-coding sequence of caspase-3^CtoG^ was PCR-amplified from *pEF-EX-BOS-Caspase-3^CtoG^-FLAG*. For *pEF-EX-BOS-7IDR^D23A^-3^CtoG^-FLAG*, the coding sequence of 7IDR^D23A^ was PCR-amplified from *pcDNA3.1-7IDR^D23A^-3-FLAG*. For *pEF-EX-BOS-7IDR^D23A, ED/A^-3^CtoG^-FLAG*, the coding sequence of 7IDR^D23A, ED/A^ was PCR-amplified from *pcDNA3.1-7IDR^D23A, ED/A^-3-FLAG*. For *pEF-EX-BOS-7IDR^Δ23^-3^CtoG^-FLAG*, the coding sequence of 7IDR^Δ23^ was PCR-amplified from *pcDNA3.1-7IDR-3-FLAG*. For *pEF-EX-BOS-7IDR^Δ23, KR/A^-3^CtoG^-FLAG*, the coding sequence of 7IDR^Δ23^ was PCR-amplified from *pcDNA3.1-7IDR^KR/A^-3-FLAG*. The two fragments (mutant 7IDR, the catalytic domain of caspase-3^CtoG^) were ligated to the *Pac*Ⅰ/*Not*Ⅰ-digested pEF-EX-BOS vector using In-Fusion.

For *pcDNA3.1-myc-mScarlet3*, the coding sequence of mScarlet3 was PCR-amplified using myc-tag harboring primers from *pmScarlet3_C1* (#189753; Addgene) and ligated to the *Eco*RⅠ*Hin*dⅢ-digested pcDNA3.1 vector using In-Fusion. For *pcDNA3.1-7IDR^D23A^-3-myc-mScarlet3*, *pcDNA3.1-7IDR^Δ23^-3-myc-mScarlet3*, and *pcDNA3.1-7IDR^Δ23, KR/A^-3-myc-mScarlet3*, each coding sequence of 7IDR-3 was PCR-amplified from *pcDNA3.1-7IDR^D23A^-3-FLAG*, respectively. The two fragments (7IDR-3 and myc-mScarlet3) were ligated to the *Eco*RⅠ/*Hin*dⅢ-digested pcDNA3.1 vector using In-Fusion.

For *pcDNA3.1-hXkr8-EGFP*, the coding sequence of hXkr8 was PCR-amplified from *pMXs-puro-hXkr8-FLAG* (a gift from S. Nagata). For *pcDNA3.1-hXkr8^2DA^-EGFP*, the coding sequence of hXkr8^2DA^ was PCR-amplified using mutation-introducing primers (D350A, GAC to GCC; D353A, GAC to GCC) from *pMXs-puro-hXkr8-FLAG*. The EGFP coding sequence was PCR-amplified from *pMXs-puro-hATP11A-EGFP*^5^. The two fragments were ligated to the *Bam*HⅠ/*Hin*dⅢ-digested pcDNA3.1 using In-Fusion. For *pcDNA3.1-Basigin-V5*, the coding sequence of Basigin was PCR-amplified using V5-tag harboring primer from *pCX4-bsr-hBasigin-cHA* (a gift from S. Nagata) and ligated to the *Bam*HⅠ/*Hin*dⅢ-digested pcDNA3.1 vector using In-Fusion. *pcDNA3.1-Basigin-V5* was co-transfected with *pcDNA3.1-hXkr8-EGFP* to enhance its localization of hXkr8-EGFP to the plasma membrane.

### Cell culture and transfection

HeLa cells (a gift from N. Mizushima) and HEK293T cells (a gift from S. Nagata) were maintained in Dulbecco’s modified Eagle’s medium (DMEM; #043-30085; FUJIFILM Wako Pure Chemical Co.) supplemented with 10% (v/v) heat-inactivated fetal bovine serum (FBS; #175012; Nichirei Biosciences) and 1% (v/v) penicillin-streptomycin (#168-23191; FUJIFILM Wako Pure Chemical Co.). MG6 cells^66,67^ (RCB2403; Riken BRC) were maintained in DMEM supplemented with 10% (v/v) heat-inactivated FBS, 1% (v/v) penicillin-streptomycin, 10 μg/mL insulin (#096-06483; FUJIFILM Wako Pure Chemical Co.), and 100 μM 2-mercaptoethanol (#137-06862; FUJIFILM Wako Pure Chemical Co.).

All cells were grown in humidified incubators, maintained at 37℃ and 5% CO_2_, and routinely assessed for mycoplasma contamination. For transfection, cells were seeded in desired plates (6-well plates [VTC-P6; Violamo]; 12-well plates [VTC-P12; Violamo]; 8-well chamber slide [SCS-N08; Matsunami]; and 35-mm glass-bottom dishes [627870; Greiner Bio-One]). After overnight incubation, cells were transfected with plasmids using PEI Max (#24765; Polysciences, Inc.) diluted in Opti-MEM (#31985070; Invitrogen). After 4 h of incubation, the medium was replaced with a growth medium. After overnight incubation, the cells were used for experiments. To inhibit apoptosis, cells were treated with 100 μM zVAD-fmk (#627610; Sigma) for 1 h before adding apoptosis-inducing reagents.

### Generation of *Caspase-3/7* KO using CRISPR-Cas9

To generate *caspase-3* KO and *caspase-7* KO HeLa cells, the cells were transfected with the *pGuide-it-tdTomato vector* (#632602; Takara) using PEI Max according to the manufacturer’s protocols. Guide RNAs were designed using CRISPRdirect^68^. The target sequence for caspase-3 (5’-GGAGAACACTGAAAACTCAG-3’) and target sequence for caspase-7 (5’-TCAGGGCTGTATTGAAGAGC-3’) were ligated to pGuide-it-tdTomato vector (linear), respectively. Twenty-four hours post-transfection, the cells were single-cell sorted into 96-well plates using the CellenOne X1i (Cellenion) single-cell isolation system with the tdTomato fluorescence signal as an indicator. Each gene deletion was validated one week after monoclonal expansion from single cells using genomic sequencing and western blotting.

### Establishment of HeLa cells stably expressing localized Dual FRET

To generate HeLa cells stably expressing localized Dual FRET probes, we used the pPiggyBac vector to avoid the recombination of mECFP and mVenus caused by viral transfection^69^. pPiggyBac vectors carrying cDNA encoding a Dual FRET system (for cytosolic, mitochondrial, ER, F-actin, or PM localization) and transposase expression vectors (*pCAGGS-PBase*, a gift from Y. Yamaguchi)^70^ were co-transfected at a ratio of 1:1 into WT, *caspase-3* KO, and *caspase-7* KO HeLa cells using PEI Max. Twenty-four hours post-transfection, cells were selected with 10 μM puromycin (#160-23151; FUJIFILM Wako Pure Chemical Co.).

### Establishment of HeLa cells stably expressing caspase-7 mutant

The retrovirus-mediated transformation was performed as described previously^63^. Briefly, the pMXs-puro vector carrying cDNA for mutant caspase variants (caspase-7, 3IDR-7, 7IDR-3, 7IDR^D23A^-3, 7IDR^ED/A^-3, or 7IDR^KR/A^-3) was transfected into HEK293T cells using FuGENE 6 (#E2691; Promega) with pEF-gag-pol-IRES-bsr^63^, pCMV-VSV-G (#RDB04392; Riken BRC), and *pAdVAntage* (#E1711; Promega). The retroviruses were produced for two days and concentrated by centrifugation at 1,500*×g* for 45 min at 4°C using Retro-X™ concentrator (#631455; Takara). After centrifugation, the supernatant was removed, and the pellet was resuspended in 500 µL of the culture medium containing 10 µg/mL polybrene (#17736-44; Nacalai Tesque). The concentrated retroviruses were then used to transform *caspase-7* KO HeLa cells, and the transformants were selected in the presence of 2 μg/mL puromycin (#19752-22; Nacalai Tesque).

### Immunocytochemistry

HeLa cells (2.5×10^3^/well) were plated in 8-well chamber slides and allowed to adhere overnight. The cells were fixed with 4% paraformaldehyde. After 15 min, cells were washed thrice in phosphate-buffered saline (PBS), followed by 15 min using 0.1% Triton X-100 in PBS (PBST) and subsequently blocked in PBST with 5% normal donkey serum (PBSTn; #S30; Millipore), and overnight incubated with primary antibodies in PBSTn at 4℃. Next, the samples were washed with PBST, incubated for 2 h with secondary antibodies in PBSTn, and washed thrice in PBST. The primary antibodies used in this study were mouse anti-ATPB monoclonal antibody (3D5; #ab14730; Abcam) and mouse anti-KDEL monoclonal antibody (#ADI-SPA-827-F; Enzo Life Sciences). The secondary antibodies used were Alexa Fluor 488-conjugated donkey anti-mouse IgG (#A-21202; Thermo Fisher Scientific) and Alexa Fluor 647-conjugated donkey anti-mouse IgG (#A-31571; Thermo Fisher Scientific). Primary and secondary antibodies were diluted to a concentration of 1:2,000. For phalloidin staining, after permeabilization with PBST, phalloidin was diluted in PBS (1:2,000 for phalloidin-488 [#A12379; ThermoFisher Scientific] and 1:2,000 for phalloidin-633 [#A22284; ThermoFisher Scientific]) and incubated for 20 min at room temperature (RT). Images were captured using a TCS SP8 microscope (Leica Microsystems).

### Live cell imaging

HeLa cells (1.0×10^5^/well) were plated in 35-mm glass-bottom dishes and allowed to adhere overnight. To avoid phenol red background, the medium was replaced with DMEM without phenol red (#040-30095; FUJIFILM Wako Pure Chemical Co.) supplemented with 10% (v/v) heat-inactivated FBS, 1% (v/v) penicillin-streptomycin, 1 mM sodium pyruvate, and 1% (v/v) Gluta-Max (#35050-061; Gibco) immediately before initiating live imaging.

Images excluding TIRF observation were captured using a Leica TCS SP8 microscope equipped with an oil-immersion objective lens (#HCX PL APO 63×/1.40-0.60 OIL CS; Leica Microsystems) and hybrid detectors (HyDs; Leica Microsystems). Each dish was placed on a stage for a 35-mm dish (#GSI-D35; Tokai Hit) maintained at 37°C under 5% CO_2_ and humidity.

mSCAT3 imaging was performed using a 448 nm diode laser and detected with HyDs, with an emission bandwidth of 450‒500 nm for mECFP and 510‒550 nm for mVenus. O-DEVD-FR imaging was performed with a 552 nm optically pumped semiconductor laser (OPSL) and detected with HyDs setting emission bandwidth of 560‒590 nm for mKOκ and 600‒670 nm for mKate2. APC-Annexin V- or HaloTag7-fused protein imaging was performed with a 638 nm diode laser and detected with HyD at a 650‒750 nm emission bandwidth. mNeonGreen or EGFP imaging was performed with a 488 nm OPSL laser and detected with HyD at an emission bandwidth of 495‒ 545 nm. mScarlet3 or pHrodo imaging was performed with a 552 nm OPSL and detected with HyD with an emission bandwidth of 560‒620 nm.

All live imaging was initiated 1 h after reagent addition and performed with only a single z-plane. Images were processed using ImageJ software (National Institutes of Health, Bethesda, MD). For apoptosis induction, 50 ng/mL TNF-α (#94948-59-1; FUJIFILM Wako Pure Chemical Co.) and 10 μg/mL CHX (#C7698; Sigma) or 1 μM staurosporine (#197-10251; FUJIFILM Wako Pure Chemical Co.) were added to the medium. To detect PS exposure during apoptosis, APC-Annexin V (1:100; #640941; BioLegend) was added to the medium.

Dual FRET images (512 × 300 pixels) were captured at 2 min intervals. Each cell was automatically segmented using the cellpose TrackMate module^71^. Because caspase activation in WT cells during apoptosis is typically completed within 40 min, we quantified the FRET ratios using 40-min images, with an approximate half-life for the mSCAT3 ratio of 20 min. However, cells expressing mutant caspase showed 1‒2 h earlier activation at the PM; therefore, the time for quantitative analysis was changed from 40 min to 1‒2 h depending on the activation time in individual cells. After segmentation using cellpose, mVenus/mECFP emission ratios for mSCAT3 and mKate2/mKOκ emission ratios for O-DEVD-FR were calculated at each time point. Next, the measured FRET ratio was interpolated to a sigmoidal curve using the “interpolate a standard curve” function and the “Sigmoidal, 4PL, X is concentration” model of GraphPad Prism 8.0 (GraphPad Software, Inc., La Jolla, CA, USA). Subsequently, the difference in the half-life of the FRET ratio (t_1/2_ (mSCAT3) - t_1/2_ (O-DEVD-FR), Δt_1/2_) was calculated. Ratiometric pseudocolor images were generated according to the macro developed by Ross *et al*.^72^ using ImageJ software. Briefly, the image stacks were split into separate stacks. Then, new images showing the FRET ratio (mVenus/mECFP or mKate2/mKOκ) were created using “Image Calculator” and converted to “physics” colored images.

APC-Annexin V (512 × 300 pixels) images were captured at 5 min intervals. hXkr8-EGFP (1024 × 600 pixels) images were captured at 10 min intervals. The mScarlet3 (512 × 300 pixels) images were captured at 2 min intervals. To quantify the time required for PS exposure and EGFP separation from the PM after caspase activation, we measured the time from the end of the decrease in mSCAT3 or O-DEVD-FR FRET ratios until each event started. HaloTag7-fused protein images (512 × 512 pixels) were captured 30 min after treatment with 100 nM HaloTag^®^ SeraFluor^TM^ 650T ligand (#A308-01; GoryoChemical). Dual FRET analysis of HeLa cells expressing HaloTag7-Lact-C2 was performed without ligands. The pHrodo (512 × 512 pixels) images were captured at 3 min intervals.

FRAP analysis was performed to compare the recovery rates of the subcellularly localized FRET probes, and cells plated in 35-mm glass-bottom dishes were transiently transfected with the desired plasmids. A 2 μm circle was photobleached (five iterations, 100%, 188 ms laser duration) in the cytoplasm using 448 nm for subcellularly localized mSCAT3 and 552 nm for subcellularly localized O-DEVD-FR. Quantification of the fluorescence recovery was performed using SP8 within a bleached region of interest.

### TIRF observation

*Caspase-7* KO HeLa cells (1.0 × 10^5^/well) were plated in 12-well plates and allowed to adhere overnight. pcDNA3.1 vectors were transfected into *caspase-7* KO HeLa cells using PEI Max. Twenty-four hours before TIRF observation, the transfected cells were seeded into the quad-free LCI-S WF chip (#LCI-SWFPQ002; Live Cell Diagnosis, Ltd.). Immediately before observation, the refractive index adjustment chamber was filled with 100% glycerol (#072-00621; FUJIFILM Wako Pure Chemical Co.), which allowed for the selective excitation of fluorophores near the glass surface by inducing total internal reflection within the glass substrate. The medium in the cell culture compartment was replaced with DMEM without phenol red.

Epifluorescence, TIRF, and bright-field images were acquired using a Nikon Eclipse Ti2 microscope equipped with a 10× dry objective lens (CFI Plan Apochromat Lambda D 10X; Nikon). The specific imaging conditions were optimized for each mode. Bright-field observations were performed by differential interference contrast (DIC) microscopy. Illumination was provided by a light-emitting diode (LED) light source, and the image contrast was enhanced using polarizers and DIC prisms. A white LED light source (Sola light engine, Lumencor) was used for epifluorescence imaging. The excitation filter was configured as a 532–552 nm band-pass filter, and a dichroic mirror transmitted wavelengths in the range of 580–650 nm. Fluorescence emission was collected through a 594–646 nm band-pass filter. For TIRF imaging, a 555 nm laser (LDI-7, 89 North) was used as the excitation source. The dichroic mirror transmitted wavelengths between 580 and 650 nm, and the emission filter was identical to that used for epifluorescence imaging (594–646 nm band-pass).

Epifluorescence and TIRF were performed to evaluate the frequency of caspase localization to the PM. After subtracting the background intensity, the TIRF/epifluorescence intensity ratio in each cell was calculated using the ImageJ software.

### Western blotting

HeLa cells (1.0 × 10^5^/well) were plated in 12-well plates containing the growth medium. After treatment, the cells were washed with PBS and lysed in RIPA buffer supplemented with an EDTA-free proteinase inhibitor (#11873580001; Roche). Protein concentrations were determined using the BCA assay (#297-73101, WAKO) following the manufacturer’s protocol. The samples were mixed with 6x Laemmli buffer, boiled at 95°C for 5 min. Each sample was separated by sodium dodecyl sulfate-polyacrylamide gel electrophoresis (SDS-PAGE), and proteins were then transferred onto Immobilon-P PVDF membranes (#IPVH00010; Millipore) for immunoblotting. The membranes were blocked with 4% skim milk (#232100; Difco Laboratories) diluted in 1×TBST. For immunoblotting, the below-listed antibodies were diluted in 4% skim milk. Signals were visualized using Immobilon Western Chemiluminescent HRP Substrate (#WBKLS0500; Millipore) and FUSION SOLO. 7S. EDGE (Vilber-Lourmat). Contrast and brightness adjustments were applied equally using the ImageJ software.

In this study, the primary antibodies used were mouse anti-V5 monoclonal antibody (#46-0705; Invitrogen), mouse anti-alpha tubulin (DM1A) monoclonal antibody (#T9026; Sigma), mouse anti-myc monoclonal antibody (#46-0603; Invitrogen), rabbit anti-Caspase-3 (EPR18297) monoclonal antibody (#ab184787; Abcam), rabbit anti-Caspase-7 polyclonal antibody (#9492; CST), and mouse anti-FLAG M2 monoclonal antibody (Sigma). The secondary antibodies used in this study were goat/rabbit/donkey anti-mouse IgG HRP-Conjugate (#W402B; Promega) and goat anti-rabbit IgG HRP-conjugated antibody (#7074S; CST). Primary and secondary antibodies were diluted to 1:5,000 and 1:10,000, respectively.

### Recombinant protein expression and purification

HEK293T cells (1.5 × 10^7^/plate) were plated in 150-mm dishes (#430599; Corning) in growth medium. After 24 h, the cells were transfected with the pEF-EX-BOS plasmid using PEI Max. For the expressed caspase variants, we introduced a mutation in the catalytic domain of the caspase (CtoG) to prevent caspases from cleaving each other during purification and adopted a cleavage-resistant form (D23A) to prevent cleavage of the IDR. Then, 20 μg of DNA per dish was mixed with 80 μg PEI Max and 2 mL Opti-MEM, incubated for 5 min at RT, and added to HEK293T cells in 15 mL Opti-MEM. After 4 h of incubation, the medium was replaced with a growth medium containing 1 mM sodium butyrate (#193-01522; FUJIFILM Wako Pure Chemical Co.). After overnight incubation, the cells were washed with PBS, detached from the substrate using trypsin, and collected. Cells were suspended in 10 mM Tris-HCl buffer (pH 7.4) containing 0.5 mM dithiothreitol (#045-08974; FUJIFILM Wako Pure Chemical Co.) and an EDTA-free protease inhibitor. The cell suspension was lysed by ultrasonication and resuspended in 10 mM Tris-HCl buffer (pH 7.4) containing 0.5 mM dithiothreitol, a protease inhibitor, 100 mM KCl, 10 mM MgCl_2_, 2 mM CaCl_2_, and 500 mM sucrose. The cell suspension was centrifuged at 8,000*×g*, 4℃ for 15 min. The supernatant was further clarified by ultracentrifugation at 100,000*×g* at 4℃ for 1 h. Then, the supernatant was mixed with 360 μL slurry (180 μL of gel) anti-FLAG magnetic beads (#M8823-1ML; Sigma) and incubated overnight at 4℃. The beads were washed with wash buffer (20 mM Tris-HCl buffer [pH 7.4] containing 0.2 mM dithiothreitol and 150 mM NaCl) and the FLAG-fused protein was eluted in wash buffer containing 150 ng/μL 3×FLAG peptide (#F4799-4MG; Sigma). The eluted proteins were purified using an Xpress Micro Dialyzer (12‒14 kDa for 7IDR-3^CtoG^-FLAG variants) or (6‒8 kDa for Lact-C2-FLAG) (#40790 and #40780; Scienova). Protein concentration was estimated by SDS-PAGE, followed by Coomassie brilliant blue staining (#178-00551; FUJIFILM Wako Pure Chemical Co.).

### Lipid overlay assay

For lipid overlay assay, 5 μL of 200 ng/μL lipid (POPS [#840034C-10MG; Sigma], POPC [#850457C-25MG; Sigma], POPE [#850757C-25MG; Sigma], POPG [#840457C-25MG; Sigma], and POPI [#850142P-100UG; Sigma]) diluted with methanol:chloroform (1:1) and spotted on supported nitrocellulose membrane (#162-0097; Bio-Rad Laboratories). Membranes were air-dried and blocked with 3% bovine serum albumin in TBST at RT for 1 h, followed by overnight incubation with 100 nM FLAG-tagged protein diluted using TBST at 4℃. Proteins were detected using mouse anti-FLAG M2 antibody (1:5,000), followed by goat/rabbit/donkey anti-mouse IgG HRP-Conjugate (1:10,000). Signals were visualized using Immobilon Western Chemiluminescent HRP Substrate and FUSION SOLO. 7S. EDGE. Contrast and brightness adjustments were applied equally using the ImageJ software.

### PS exposure assay

HeLa cells (2.0 × 10^5^/well) were plated in 6-well plates in a growth medium. 6 h post-treatment with TNF-α and CHX, cells were collected and washed with PBS and resuspended in 400 μL Annexin V binding buffer (#422201; BioLegend). Next, the cells were mixed with 200 μL Annexin V binding buffer containing 500 nM CellEvent caspase-3/7 Green (#C10427; Invitrogen). 30 min after incubation at RT, the cell suspension was mixed with APC-Annexin V (#640941; BioLegend) (1:20). After 15 min of incubation at RT, the cells were washed twice with Annexin V binding buffer. The cells were then analyzed using a FACSAria III (BD Biosciences).

### Efferocytosis assay

For flow cytometry analysis, MG6 cells (4.0 × 10^5^/well) were plated in 12-well plates in a growth medium containing 1% methylcellulose (#M0387-100G; Sigma), centrifuged at 500*×g* for 2 min at RT using PlateSpinⅡ (KUBOTA), and overnight incubated at 37℃. To prepare apoptotic HeLa cells, cells plated in 100-mm dishes (#430167; Corning) were treated with TNF-α (50 ng/mL) and CHX (10 μg/mL). 6 h post-treatment, all HeLa cells, including surviving cells, were collected, washed with PBS, and stained with 150 nM pHrodo Red succinimidyl ester (#p36600; Invitrogen) for 20 min at RT. Then, pHrodo-labeled HeLa cells were added to MG6 cells in a growth medium containing 1% methylcellulose (MG6:HeLa = 2:1), followed by centrifugation at 500*×g* for 2 min at RT. Cells were collected 0, 20, 40, 60, 120 min after co-culture, blocked with 2% FBS, and stained with 200 ng/µL APC-anti-CD11b (#101212; BioLegend) for 15 min at RT. The cells were then suspended in 20 mM CHES-NaOH buffer (pH 9.0) containing 150 mM NaCl and 2% FBS and analyzed using a FACSAria IIIu (BD Biosciences).

For live imaging, MG6 cells (1.0 × 10^5^/well) were plated in 35-mm glass-bottom dishes in a growth medium containing 1% methylcellulose, centrifuged at 500 ×g for 2 min at RT using PlateSpinⅡ and incubated overnight at 37℃. pHrodo-labeled HeLa cells were then added to the MG6 cells (MG6: HeLa = 10:1) followed by centrifugation at 500 ×g for 2 min at RT. The images were captured using a TCS SP8 microscope (Leica Biosciences).

### IDR prediction

The amino acid sequences obtained from the GenBank database and the NCBI Reference Sequence database are as follows: Human caspase-3 (GenBank CAC88866.1), human caspase-7 (NCBI RefSeq NP_001253985.1), *Mus musculus* caspase-7 (NCBI RefSeq NP_031637.1), *Lupus europaeus* caspase-7 (NCBI RefSeq XP_062069875.1), *Camelus ferus* caspase-7 (NCBI RefSeq XP_014413725.1), *Molossus molossus* caspase-7 isoform X1 (NCBI RefSeq XP_036121468.1), *Gallus gallus* caspase-7 isoform X2 (NCBI RefSeq XP_421764.3), *Python bivittatus* caspase-7 (NCBI RefSeq XP_007441954.1), *Trachemys scripta elegans* caspase-7 (NCBI RefSeq XP_034632910.1), *Crocodylus porosus* caspase-7 isoform X2 (NCBI RefSeq XP_019411760.1), *Xenopus laevis* caspase-7 L homolog (NCBI RefSeq NP_001081408.1), *Cynops orientalis* caspase-7 (GenBank AFN55259.1), *Danio rerio* caspase-7 (NCBI RefSeq NP_001018443.2), *Heterodontus francisci* caspase-7 (NCBI RefSeq XP_067909046.1), *Branchiostoma floridae* caspase-7-like isoform X1 (NCBI RefSeq XP_035669585.1), *Ciona intestinalis* caspase-7 (NCBI RefSeq XP_018669775.2), Drice (NCBI RefSeq NP_524551.2), Dcp-1 (NCBI RefSeq NP_476974.1), and *Hydra vulgaris* caspase-7 (NCBI RefSeq NP_001267753.1). The IDR of each protein was predicted using the PSIPRED protein sequence analysis workbench in DISOPRED3 (http://bioinf.cs.ucl.ac.uk/psipred/). The highly basic and hydrophobic regions of caspase-3 and caspase-7 were predicted using the BH search tool^34^ (http://helixweb.nih.gov/bhsearch).

### Statistical analysis

Statistical analyses were performed using GraphPad Prism 8.0 (GraphPad Software, Inc). The dotted plots show the means ± standard error of the mean (SEM). Time courses show the means ± standard deviation (SD). *P*-values were calculated using one-way analysis of variance (ANOVA) with Dunnett’s comparison test or unpaired two-tailed *t*-test. NS: *P* > 0.05; *: *P* < 0.05; **: *P* < 0.01; ***: *P* < 0.001; ****: *P* < 0.0001. Flow cytometry data were analyzed using FlowJo version 10.10.0 (Becton Dickinson).

## Acknowledgments

We thank Dr. H. Imamura, Dr. A. Koto, Dr. T. Kitamura, Dr. S. Nagata, Dr. Y. Yamaguchi, and Dr. H. Miyoshi for providing plasmids. We thank Dr. N. Mizushima and Dr. S. Nagata for providing cell lines. We thank Dr. Y. Kishi, Dr. A. Watanabe, and the One-Stop Sharing Faculty Center for Future Drug Discovery at the Graduate School of Pharmaceutical Sciences, University of Tokyo, for the FACS analysis. We thank all members of Miura Lab for their technical assistance and discussions, with special gratitude to R. Takamoto for providing experimental support. We thank *Editage* (www.editage.com) for the English language editing of this manuscript. This work was supported by grants from the Ministry of Education, Culture, Sports, Science, and Technology of Japan (KAKENHI Grant Numbers 21K15080, 22H05586, and 23K05747 to N.S. and 21H04774, 23H04766, and 24H00567 to M.M.); the Japan Agency for Medical Research and Development (AMED; Grant number JP21gm5010001 to M.M.); and the Takeda Science Foundation and Sumitomo Foundation to N.S. Y.T. is a research fellow of the Japan Society for the Promotion of Science (KAKENHI Grant 24KJ0804).

## Author contributions

Author contributions: Y.T., M.M., and N.S. designed the study; Y.T. conducted most of the experiments and analyzed most of the data under the supervision of M.M. and N.S.; Y.M. and K.S. established stable cell lines and assisted with the lipid overlay assay; Y.Z. and Y.S. performed the TIRF analysis; Y.T., M.M., and N.S. wrote the manuscript.

## Competing interest declaration

The authors declare that they have no conflict of interest.

---

**Extended Data Figure 1.**
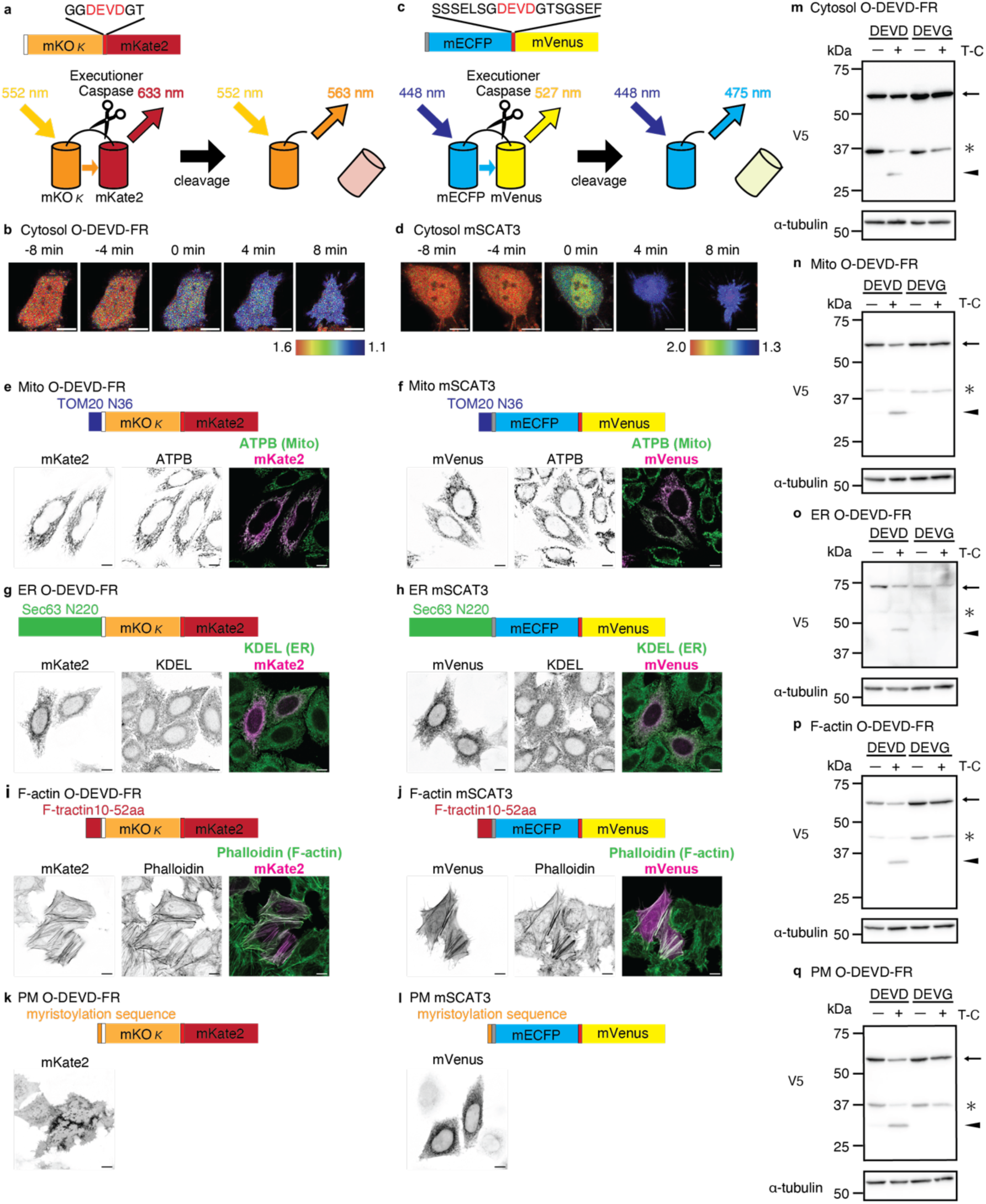
Characterization of subcellularly localized FRET biosensors. (a, c) Schematic representation depicting FRET biosensors developed to detect executioner caspase activation. O-DEVD-FR comprises mKOκ, mKate2, and V5-tag (white box) fused to the N-terminus of O-DEVD-FR (a). mSCAT3 comprises mECFP, mVenus, and myc-tag (gray box) fused to the N-terminus of mSCAT3 (c). (b, d) FRET ratio changes in cytosolic O-DEVD-FR and mSCAT3 during extrinsic apoptosis at the indicated time. Zero min indicates the time at which the FRET ratio reaches half the maximum and minimum values. Scale bars, 10 μm (e, f) Representative images of mitochondria (Mito)-localized O-DEVD-FR and mSCAT3 in HeLa cells. Scale bars, 10 μm (g, h) Representative images of ER-localized O-DEVD-FR and mSCAT3 in HeLa cells. Scale bars, 10 μm (i, j) Representative images of F-actin-localized O-DEVD-FR and mSCAT3 in HeLa cells. Scale bars, 10 μm (k, l) Representative images of the plasma membrane (PM)-localized O-DEVD-FR and mSCAT3 in HeLa cells. Scale bars, 10 μm. (m-q) Western blot of O-DEVD-FR transiently expressed in HeLa cells exposed to TNF-α/CHX (T-C). (m, Cytosol; n, Mito; o, ER; p, F-actin; q, PM). Black arrows indicate full-length localized O-DEVD-FR (Expected Mw: m, 52.4 kDa; n, 56.3 kDa; o, 77.6 kDa; p, 56.6 kDa; q, 53.2 kDa). Black arrowheads indicate cleaved localized O-DEVD-FR (Expected Mw: m, 26.4 kDa; n, 30.3 kDa; o, 51.6 kDa; p, 30.6 kDa; q, 27.2 kDa). Asterisks indicate the non-specific cleavage products of O-DEVD-FR.

**Extended Data Figure 2.**
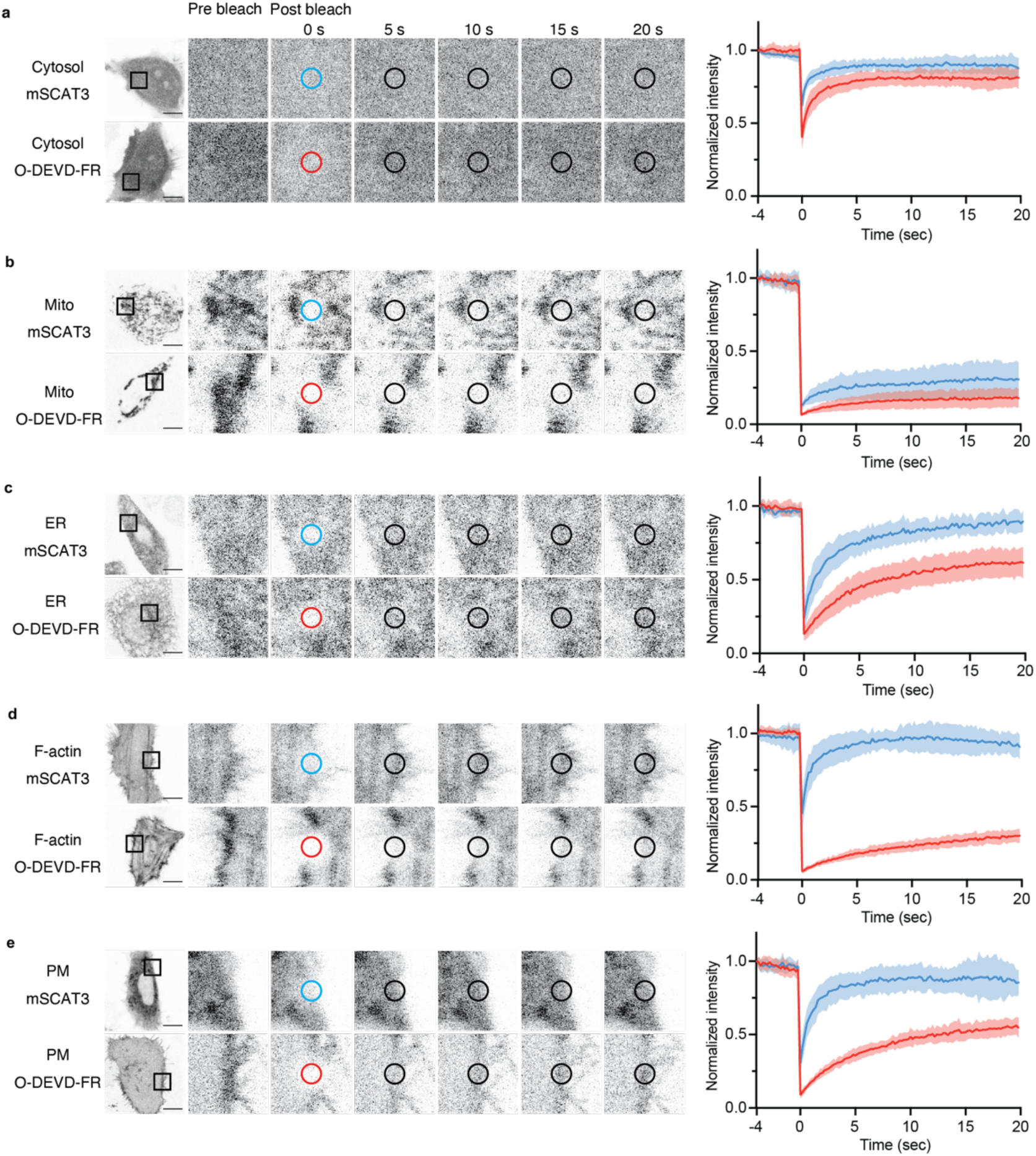
O-DEVD-FR stably localizes to subcellular regions compared with mSCAT3. (a-e) FRAP analysis of subcellularly localized mSCAT3 and O-DEVD-FR. ((a) Cytosol; (b) mitochondria (Mito); (c) ER; (d) F-actin; (e) plasma membrane (PM)). In all the figures, circles represent the point of photobleaching. Fluorescence recovery at the bleached point was measured. Zero sec indicate the time at which the circular area was photobleached. The fluorescence intensity is normalized by dividing the value at t = -4 sec. All the molecules were transiently expressed in WT HeLa cells. Solid lines indicate the mean, and shaded areas indicate SD. n =10 cells per experiment. Red represents O-DEVD-FR, and blue represents mSCAT3. Scale bars, 10 μm.

**Extended Data Figure 3.**
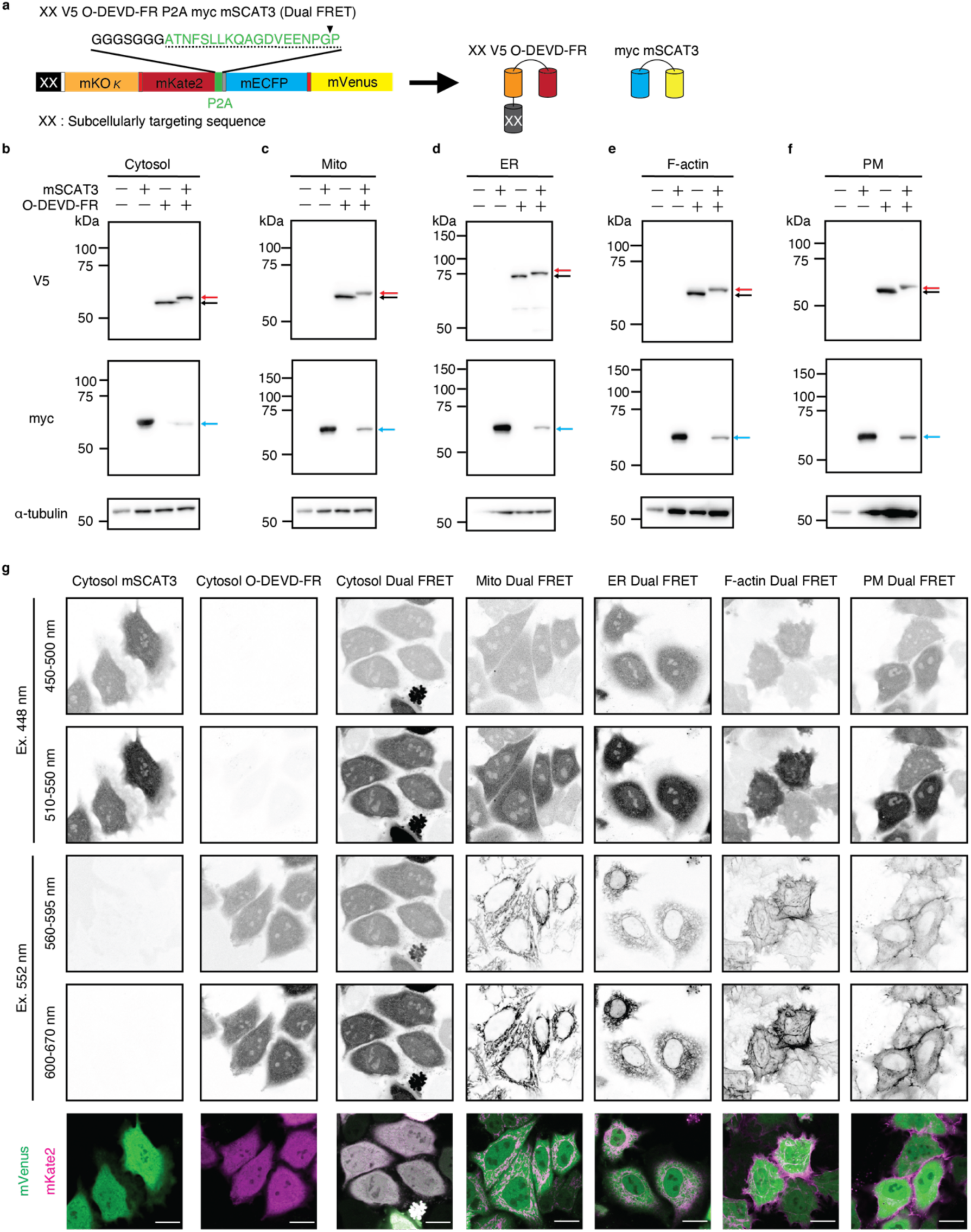
Characterization of subcellularly targeted O-DEVD-FR-P2A-cytosolic mSCAT3. (a) A schematic diagram depicting the subcellular localization of V5-O-DEVD-FR-P2A-myc-mSCAT3 (Dual FRET). In Dual FRET, a flexible GGGSGGG linker and a P2A peptide (dotted line) were inserted between O-DEVD-FR and mSCAT3^DEVD^. The black arrowhead indicates the P2A peptide separation site. XX indicates the subcellularly targeting sequence. (b-f) Western blot of mSCAT3^DEVD^, subcellularly localized O-DEVD-FR, or Dual FRET expressing HeLa cells. Black arrows indicate full-length O-DEVD-FR. Red arrows indicate the full-length localized O-DEVD-FR fused to the P2A peptide. Blue arrows indicate full-length myc mSCAT3. (g) Representative images of mSCAT3, O-DEVD-FR, and Dual FRET transiently expressed in HeLa cells. Scale bars, 10 μm. For all figures, all molecules were transiently expressed in WT HeLa cells.

**Extended Data Figure 4.**
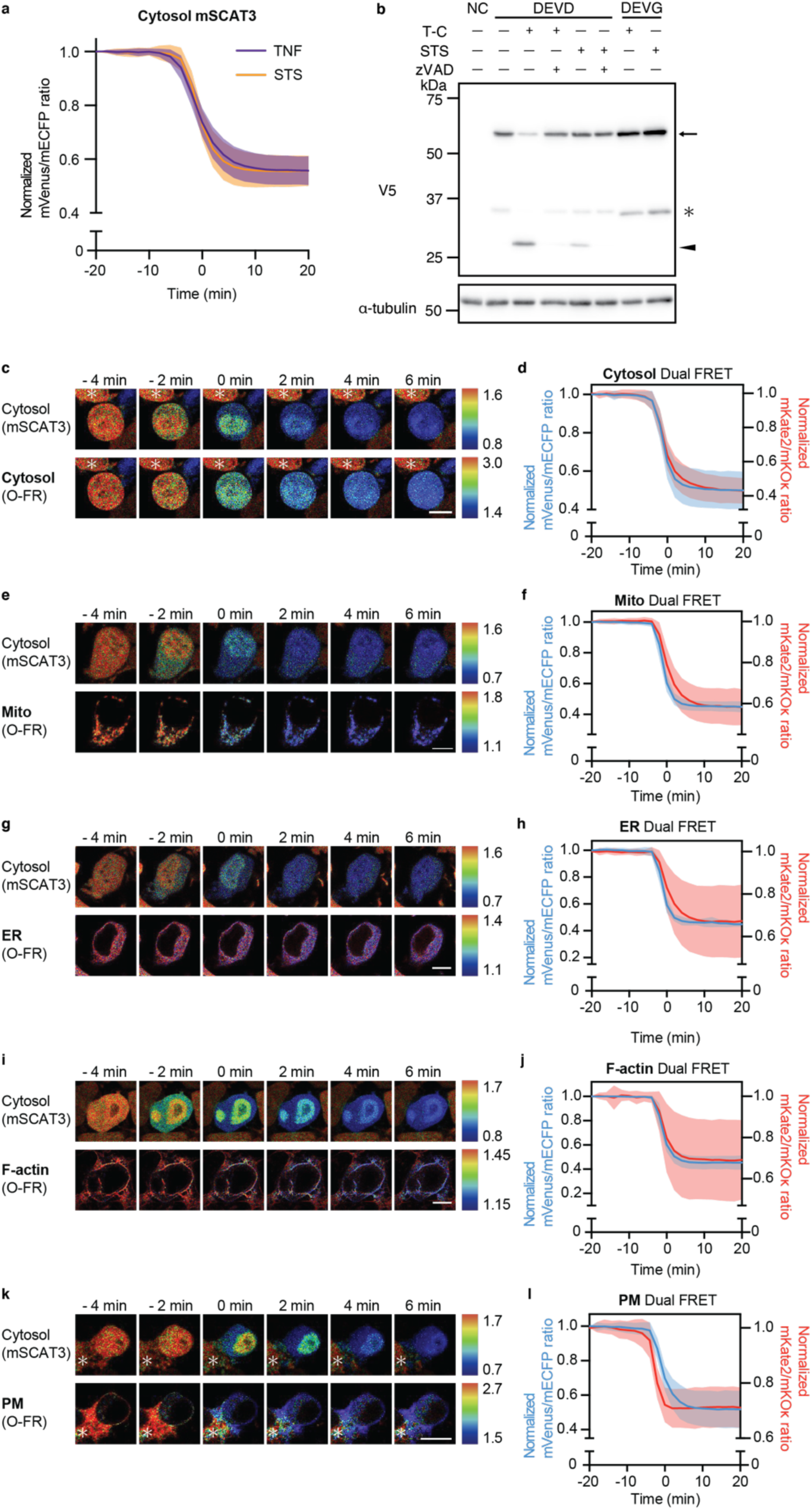
Subcellularly targeted caspase activity during intrinsic apoptosis. (a) Average cytosolic caspase activity in WT HeLa cells during extrinsic (purple) and intrinsic apoptosis (orange). The FRET ratio is normalized by dividing the ratio at t = -20 min. (b) Western blot of V5-O-DEVD-FR cleavage dependent on TNF-α/CHX or staurosporine-induced apoptosis. To confirm the dependence on caspase activity, zVAD was added 1 h before adding the apoptosis-inducing reagent. The black arrow indicates full-length O-DEVD-FR. The black arrowhead indicates cleaved O-DEVD-FR. The asterisk indicates non-specific cleavage products of O-DEVD-FR. (c-l) Live imaging analysis of Dual FRET stably expressed in WT HeLa cells during intrinsic apoptosis at the indicated times. Zero min indicates the time at which the mVenus/mECFP ratio reaches half the maximum and minimum values. The FRET ratio is normalized by dividing the ratio at t = -20 min. (c, e, g, i, k) Representative pseudo-colored FRET ratio images of cytosolic mSCAT3 (top: mVenus/mECFP) and subcellularly localized O-DEVD-FR (bottom: mKate2/mKOκ) in HeLa cells during intrinsic apoptosis. ((c) Cytosol, (e) mitochondria (Mito), (g) ER, (i) F-actin, and (k) plasma membrane (PM)). (d, f, h, j, l) Average response of Dual FRET stably expressed in WT HeLa cells during extrinsic apoptosis (blue, cytosolic mSCAT3 vs. red, subcellular O-DEVD-FR). ((d) Cytosol, n = 45; (f) Mito, n = 31; (h) ER, n = 23; (j) F-actin, n = 33; (l) PM, n = 40 cells). Asterisks indicate neighboring cells. Scale bars, 10 μm. For all figures, time courses show the mean ± SD.

**Extended Data Figure 5.**
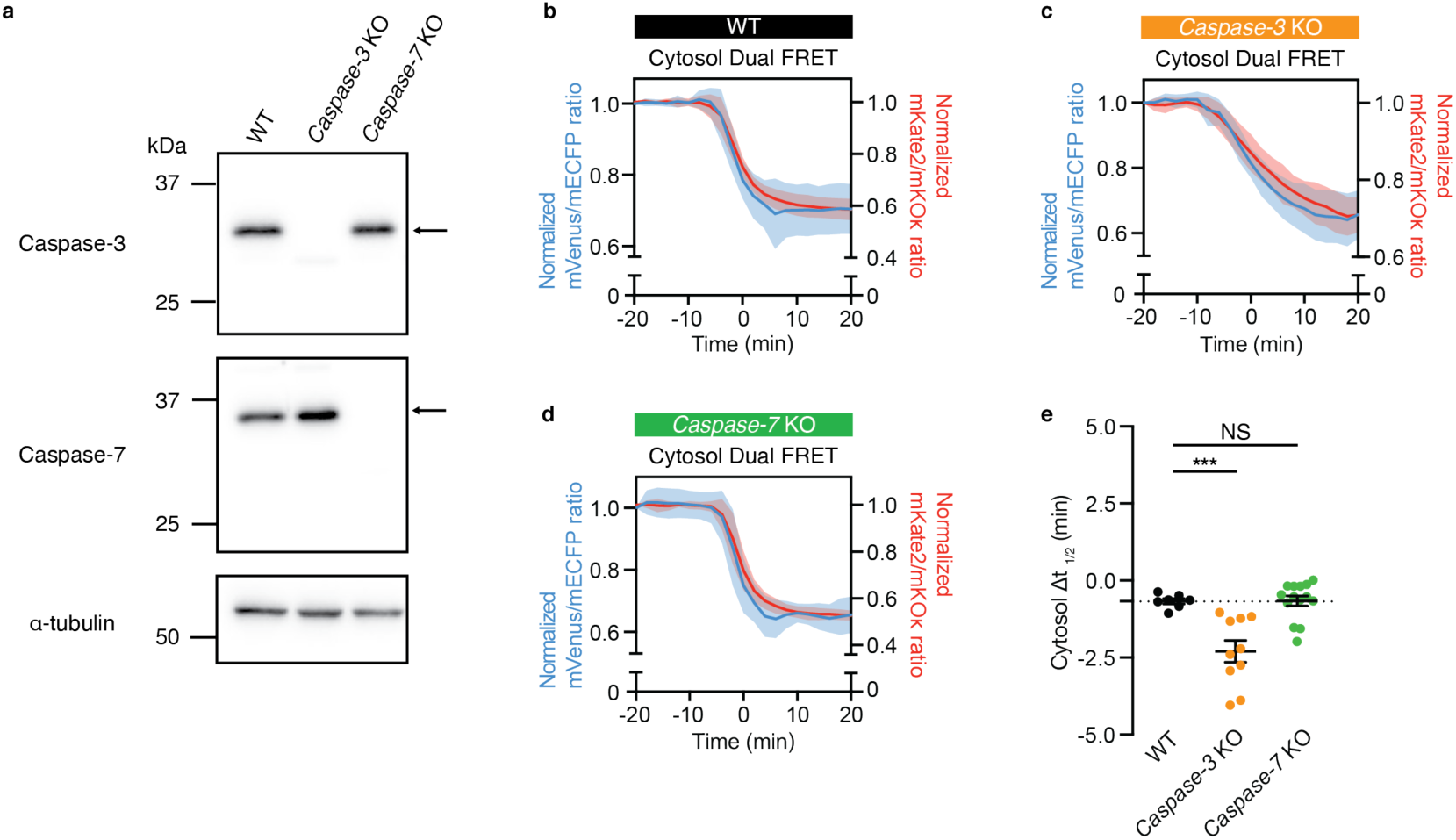
Establishment of caspase-3 KO and caspase-7 KO HeLa cells. (a) Western blot of WT, *caspase-3* KO, and *caspase-7* KO HeLa cells. (b-d) Average response of Cytosol Dual FRET in WT, *caspase-3* KO, and *caspase-7* KO HeLa cells during extrinsic apoptosis (blue, cytosolic mSCAT3 vs. red, cytosolic O-DEVD-FR). The FRET ratio is normalized by dividing the ratio at t = -20 min. (e) Δt_1/2_ of Cytosol Dual FRET in WT (black, n = 7 cells), *caspase-3* KO (orange, n = 10 cells), and *caspase-7* KO HeLa (green, n = 16 cells) during extrinsic apoptosis. Statistical analysis was performed using one-way ANOVA with Dunnett’s comparison test. NS: *P* > 0.05; ***: *P* < 0.001.

**Extended Data Figure 6.**
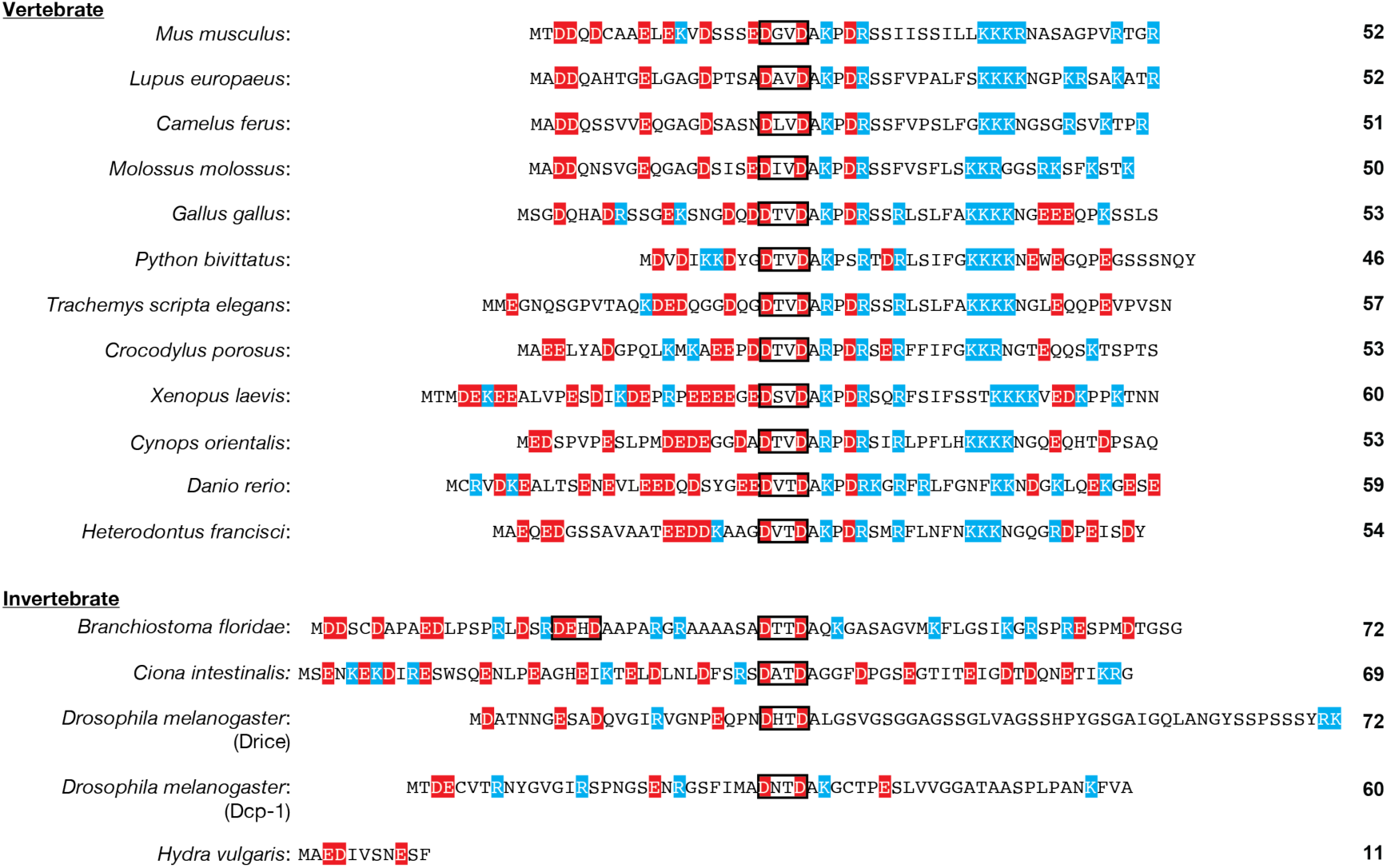
Alignment of amino acid sequences of caspase-7 IDR derived from other animal species. Alignment of amino acid sequences of caspase-7 IDR derived from vertebrate caspase-7 orthologs, invertebrate caspase-7 orthologs, and two types of *Drosophila* executioner caspases. Acidic amino acids are indicated in red. Basic amino acids are shown in blue. The predicted caspase recognition sites are shown in boxes.

**Extended Data Figure 7.**
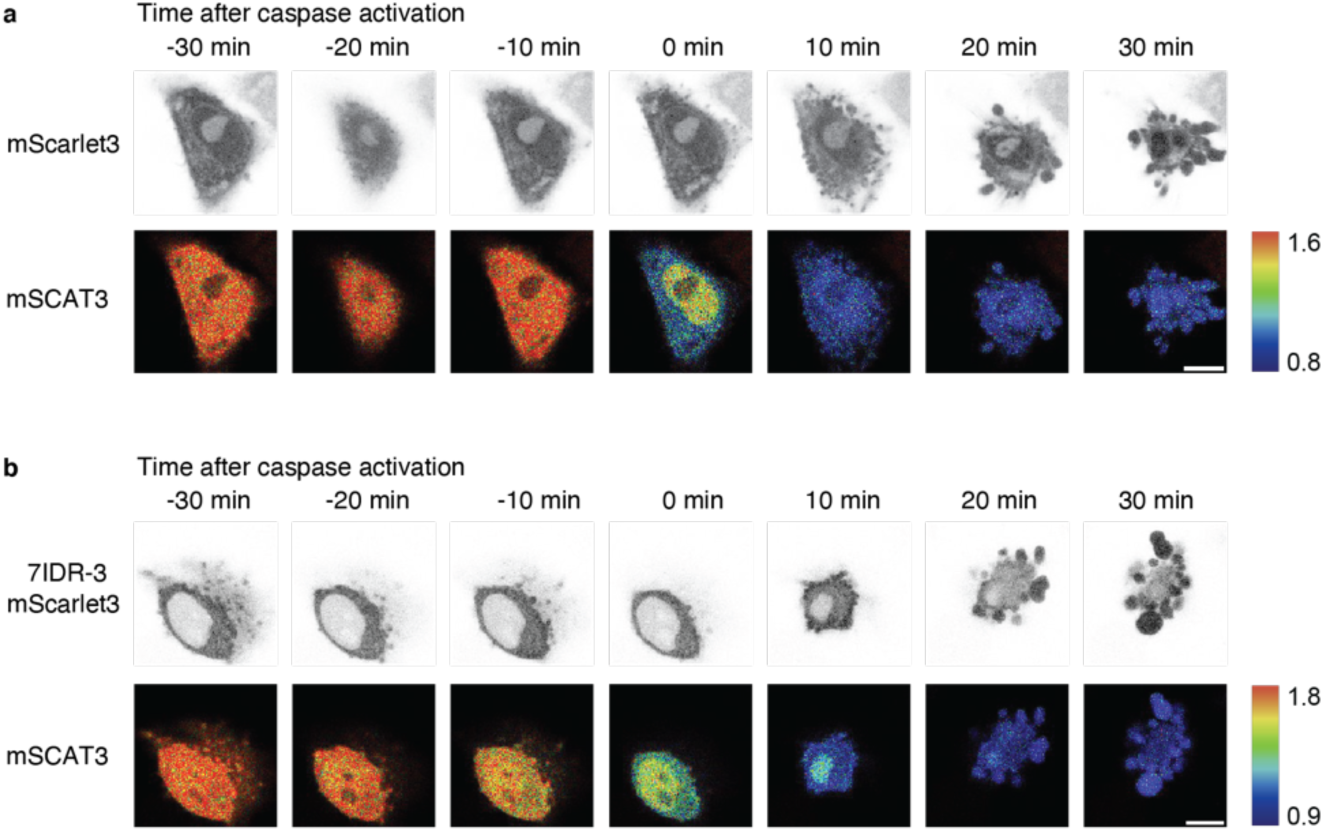
Caspase localization dynamics during extrinsic apoptosis based on confocal microscopy. (a, b) Live imaging of mScarlet3 and mSCAT3 transiently expressed in *caspase-7* KO HeLa cells during extrinsic apoptosis at the indicated time ((a), mScarlet3; (b) 7IDR-3 C-terminally fused with mScarlet3). Zero min indicates the time at which the mVenus/mECFP ratio reaches half the maximum and minimum. Representative images of mScarlet3 (top) and pseudo-colored FRET ratio images of cytosolic mSCAT3 (bottom: mVenus/mECFP) in HeLa cells during extrinsic apoptosis. Scale bars, 10 μm.

**Extended Data Figure 8.**
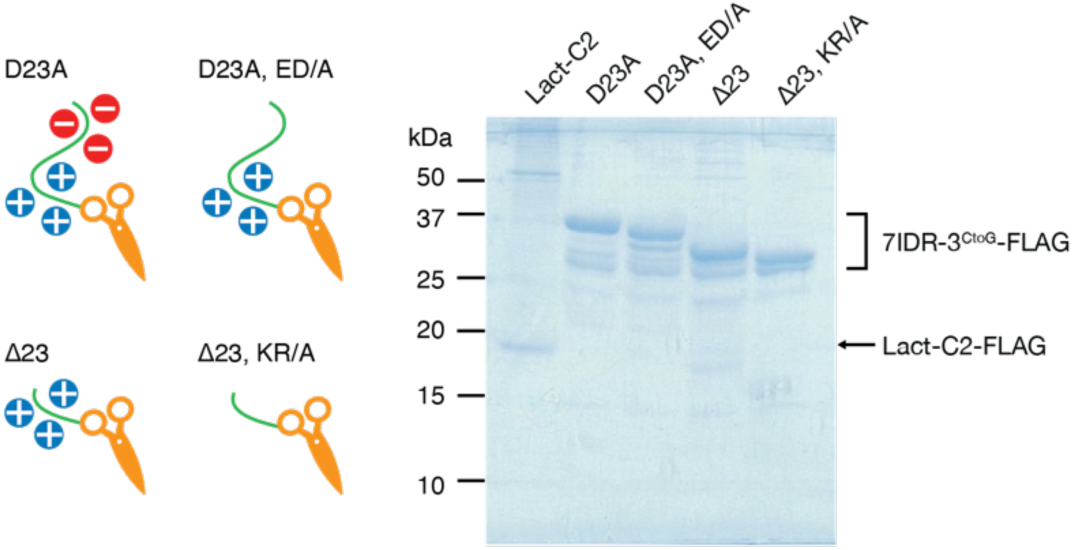
Coomassie brilliant blue staining of recombinant proteins. Coomassie brilliant blue staining of recombinant Lact-C2-FLAG (19.2 kDa), 7IDR^D23A^-3^CtoG^-FLAG (35.2 kDa), 7IDR^D23A, ED/A^-3^CtoG^-FLAG (34.8 kDa), 7IDR^Δ23^-3^CtoG^-FLAG (32.9 kDa), and 7IDR^Δ23, KR/A^-3^CtoG^-FLAG (32.3 kDa) purified from HEK293T cells.

**Extended Data Figure 9.**
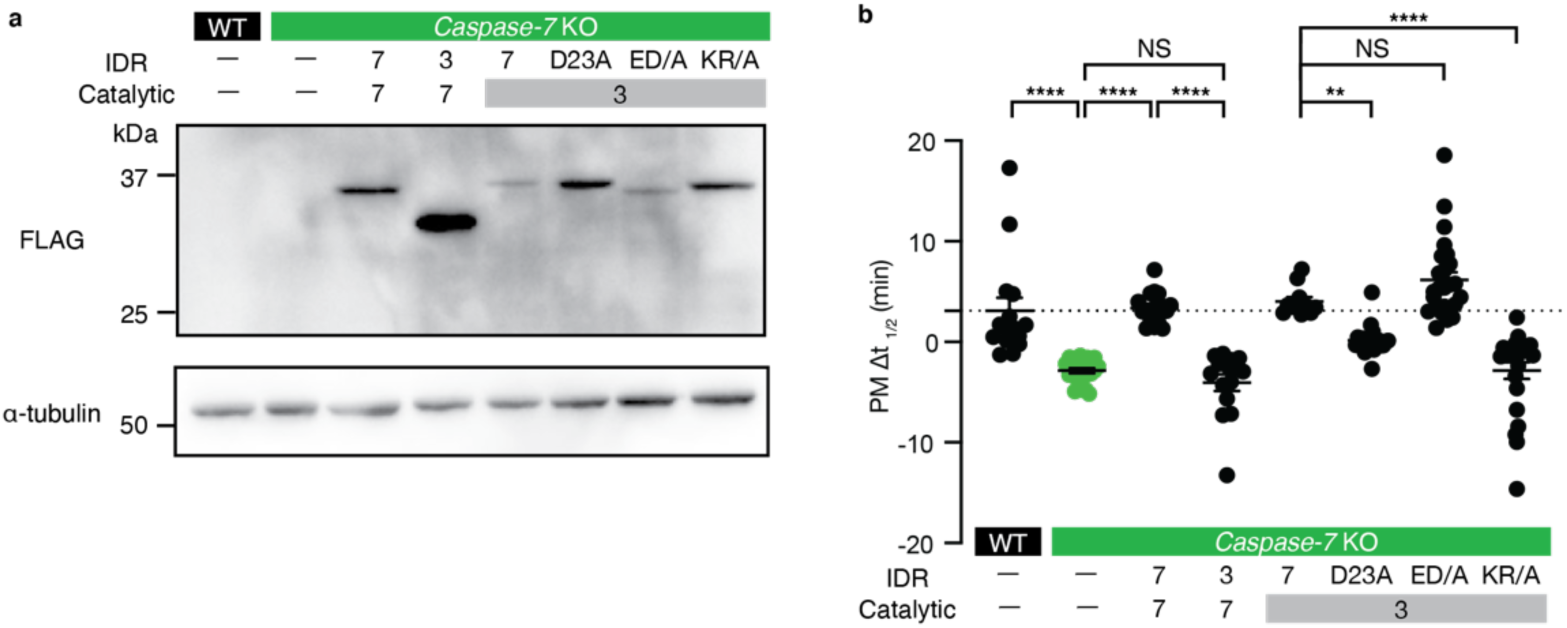
Establishment of HeLa cells stably expressing mutant caspase via retroviral transduction. (a) Western blot of WT, *caspase-7* KO, and *caspase-7* KO HeLa cells stably expressing caspase-7-FLAG, 3IDR-7-FLAG, 7IDR-3-FLAG, 7IDR^D23A^-3-FLAG, 7IDR^ED/A^-3-FLAG, and 7IDR^KR/A^-3-FLAG. (b) Δt_1/2_ of PM Dual FRET transiently expressed in WT (n = 15 cells), *caspase-7* KO (n = 23 cells), and *caspase-7* KO HeLa cells stably expressing caspase-7-FLAG (n = 18 cells), 3IDR-7-FLAG (n = 11 cells), 7IDR-3-FLAG (n = 15 cells), 7IDR^D23A^-3-FLAG (n = 19 cells), 7IDR^ED/A^-3-FLAG (n = 25 cells), and 7IDR^KR/A^-3-FLAG (n = 24 cells) HeLa cells during extrinsic apoptosis. For all figures, dot plots show the mean ± SEM. Statistical analysis was performed using one-way ANOVA with Dunnett’s multiple comparison test. NS: *P* > 0.05; **: *P* < 0.01; ****: *P* < 0.0001.

**Extended Data Figure 10.**
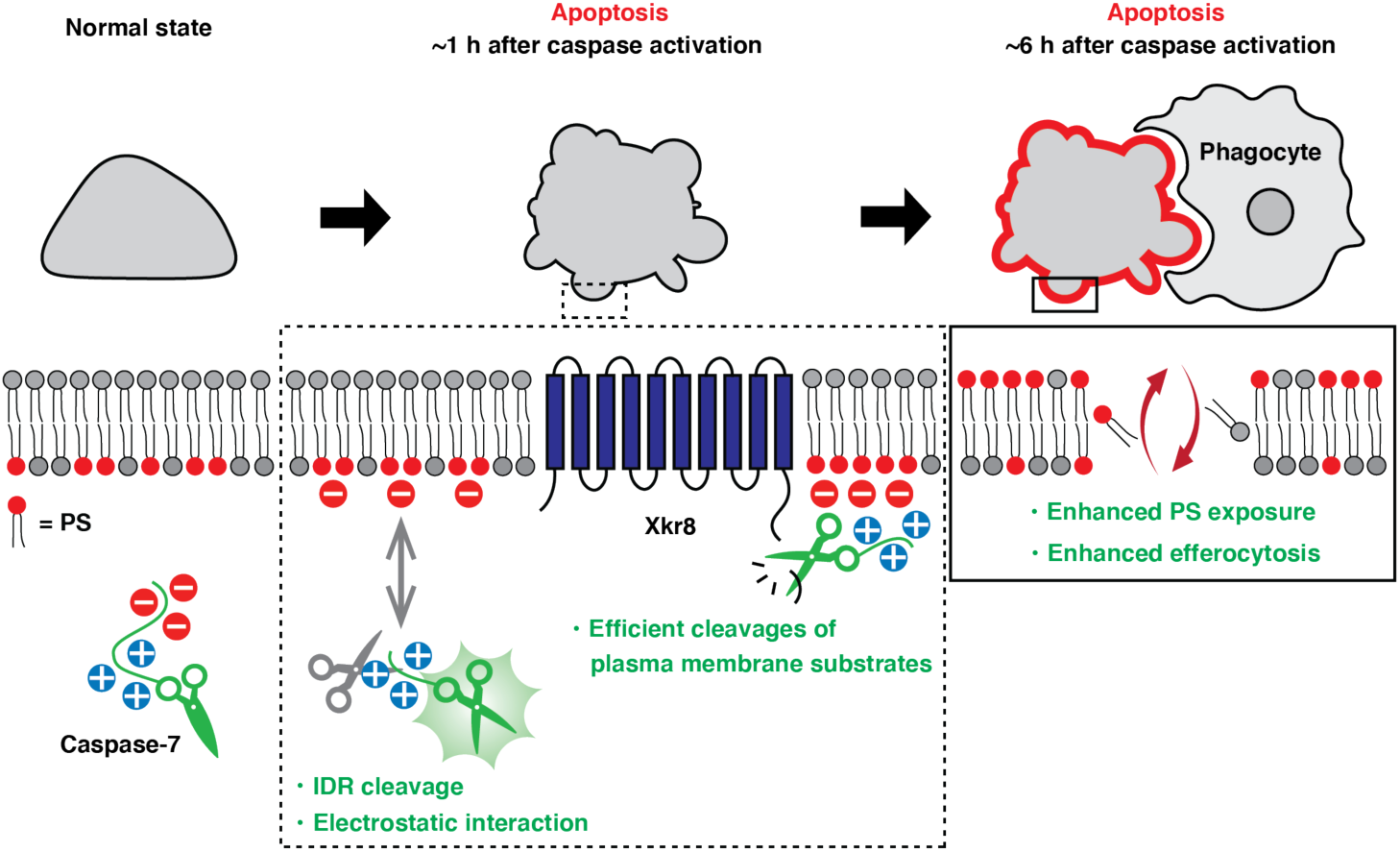
Proposed model. Under normal conditions, the executioner caspases are inactive zymogens within the cell. The executioner caspase is activated in an apoptotic stimuli-dependent manner, and the prodomain of caspase-7 is cleaved, resulting in the separation of the polybasic domain from the polyacidic domain. This allows the polybasic domain to electrostatically interact with PS on the plasma membrane (PM), resulting in early peak activation at the PM. Activated caspase-7 then cleaves and activates scramblase Xkr8, thereby promoting rapid PS exposure and subsequent efferocytosis.

## Notes

### Competing Interest Statement

The authors have declared no competing interest.

